# Integrated single cell multiomic profiling and functional validation reveal distinct cellular routes to human plasma cell differentiation

**DOI:** 10.64898/2026.02.16.706247

**Authors:** Colin A. Fields, James F. Read, Heather Coffman, Edward P. Petrow, Anthony Bosco, Deepta Bhattacharya

## Abstract

Upon activation, B cells undergo profound transcriptional, epigenetic, and metabolic reprogramming to form antibody secreting plasma cells bearing little resemblance to their progenitors. Here, we used single cell RNA and ATAC sequencing of primary and *in vitro* differentiating human B cells to identify multiple distinct plasma cell subsets and differentiation routes depending on the starting cell type. In primary tonsils, we observed two distinct plasma cell subsets distinguished by expression of CD44 variant 9 (CD44v9), CD38, CD31, and CD10. A transient and rare CD30^+^ intermediate was identified both in primary tonsils and *in vitro*. CD40, TLR9, and cytokine stimulation of naïve and memory B cells yielded CD30+ cells, which then formed plasma cells that were exclusively CD44v9^+^. CD30^+^ intermediates were not generated from primary germinal center B cells as they differentiated into plasma cells, which mostly lacked expression of CD44v9. Analysis of single cell multiomic data and pharmacological inhibition experiments demonstrated that the CD30^+^ intermediate was promoted by the transcription factor MEF2C. BAFF and APRIL promoted further maturation of these cells to CD44v9^+^ plasma cells. These data suggest that human germinal center-independent and -dependent ontogenies are biased towards distinct differentiation routes and terminal plasma cells.

## Introduction

Antibody-secreting plasma cells are generated upon activation and differentiation of upstream B cells. Plasma cells can be generated from B cells through a variety of activation cues, such as crosslinking of the B cell receptor (BCR), signaling through CD40, engagement of Toll-like receptors, and instructive cytokines (Jego et al., 2003; Kräutler et al., 2017a; Poeck et al., 2004). These signals integrate to induce dramatic changes, resulting in terminally differentiated plasma cells bearing little resemblance to other B-lineage cells. Irrespective of the activation cues, a common core set of programs changes as B cells differentiate towards plasma cells. Paired Box 5 (PAX5)-driven B cell transcriptional and epigenetic programs are extinguished by PR/SET Domain 1 (Blimp-1) (Lin et al., 2002), which works in concert with Interferon Regulatory Factor 4 (IRF4) to enforce the plasma cell program (Sciammas et al., 2006). As the cells differentiate, they begin secreting vast quantities of antibodies, leading to activation of X-Box Binding Protein 1 (XBP1) in response to the resulting ER stress (Gass et al., 2002). To cope with the bioenergetic demands of antibody secretion, they also undergo metabolic reprogramming driven by Protein Kinase C (PKC) and Mammalian Target of Rapamycin Complex 1 (mTORC1) signaling (Tsui et al., 2018).

The studies that revealed these mechanistic details were first mostly conducted *in vitro*, as the cellular intermediates between B cells and plasma cells are exceedingly rare *in vivo*. Perhaps these cellular intermediates are particularly transient *in vivo*, with transition states progressing rapidly and abruptly. While mouse studies have employed various approaches, such as genetic reporters, to identify these rare intermediates (Gómez-Escolar et al., 2022; Ise et al., 2018; Kräutler et al., 2017b; Schulz et al., 2025), this experimental strategy is not feasible for human B cells, which differ from mouse B cells in fundamentally important ways, such as the expression of innate pattern recognition receptors (Bekeredjian-Ding and Jego, 2009). Several studies have thus employed single cell transcriptional and epigenetic profiling to infer developmental trajectories of differentiating human B cells (Alaterre et al., 2024; Verstegen et al., 2023). Functional validation of the predictions made by these studies is a critical next step given that dimensionality reduction of single cell data can misrepresent developmental relationships (Alquicira-Hernandez et al., 2021; Chari and Pachter, 2023; Wilk et al., 2020). Coupling these types of single cell profiling studies with functional validation and perturbation of cellular intermediates, transcriptional programs, and signaling pathways could provide further insight into how human plasma cells develop. Despite these advances in understanding plasma cell biology, critical questions remain unanswered for human systems. First, do distinct activation pathways lead to functionally divergent plasma cell fates? Second, what are the transcriptional regulators that could be therapeutically targeted to enhance plasma cell generation? Third, can we identify rare cellular intermediates that represent decision points in cell fate commitment? Answering these questions requires integrating transcriptional and epigenetic profiling with functional perturbation studies.

While all plasma cells start out as B cells, there are multiple routes they can take to become a plasma cell (Eisenbarth et al., 2025). These pathways to plasma cells can be broken down into germinal center-dependent (GC-dependent) or GC-independent pathways. GC-dependent pathways involve a germinal center B cell intermediate which traffics back and forth between dark and light zones, accumulating somatic hypermutations and eventually terminally differentiating into plasma cells (Young and Brink, 2021). GC-independent pathways allow B cells to “directly” differentiate into plasma cells (Bortnick and Allman, 2013). Additionally, plasma cells can take up residence in various locations within the body. The bone marrow is somewhat unique in containing repertoire of plasma cells generated from a wide variety of conditions (Ferreira-Gomes et al., 2024; Lemke et al., 2016; Tellier et al., 2024a; Wilmore et al., 2021, 2018).

Here, we integrated multiomic profiling of primary and in vitro differentiating human B cells with systematic functional validation to define the cellular routes and molecular programs governing plasma cell differentiation. We identified two distinct plasma cell populations distinguished by CD44v9 expression. Through gene regulatory network inference and targeted perturbation experiments, we discovered that MEF2C, STAT1, POU2F2, XBP1, and IRF4 collectively regulate a rare CD30+ intermediate that serves as a gateway to CD44v9+ plasma cells. Pharmacological modulation of these factors revealed opportunities to enhance plasma cell yields, with implications for vaccine development and therapeutic antibody production

## Results

### *In vitro* differentiation of primary B cells fills gaps in the plasma differentiation trajectory

To epigenetically and transcriptionally profile the starting and ending cell types during plasma cell differentiation, we performed single cell paired ATAC and RNA sequencing on CD19^+^ human primary tonsil B-lineage cells sorted according to the scheme depicted in (Fig 1A) with naïve B cells defined as CD27^-^CD38^-^, memory as CD27^+^CD38^-^, germinal center as CD27^+/-^CD38^+^, and plasma cells as CD27^++^CD38^++^. Data were embedded in a UMAP space based on integration of both ATAC and transcriptional profiles of these B lineage cells (Fig 1B). Cell identities were assigned based on expression levels of known marker genes, such as *PAX5, BCL6, PRDM1,* antibody isotype, and cell cycle genes (*MKI67*) as depicted in (Fig 1C). This allowed for the identification of memory B cell subsets, light zone (non-cell cycling) or dark zone (cell cycling) germinal center cells, and two separate clusters of plasma cells. Yet very few cells were identified in the space between clusters, likely due to rapid terminal differentiation *in vivo* after cells pass a critical threshold for plasma cell differentiation, precluding inference of developmental trajectories to either plasma cell subset. This prompted us to perform the same sequencing and embedding on *in vitro* differentiating naïve B cells. The main advantages of this system are the ability to have many cells differentiating in synchrony and to sample at very early stages post-activation, allowing for the identification of populations that are rare or very transient *in vivo*.

**Fig. 1.**
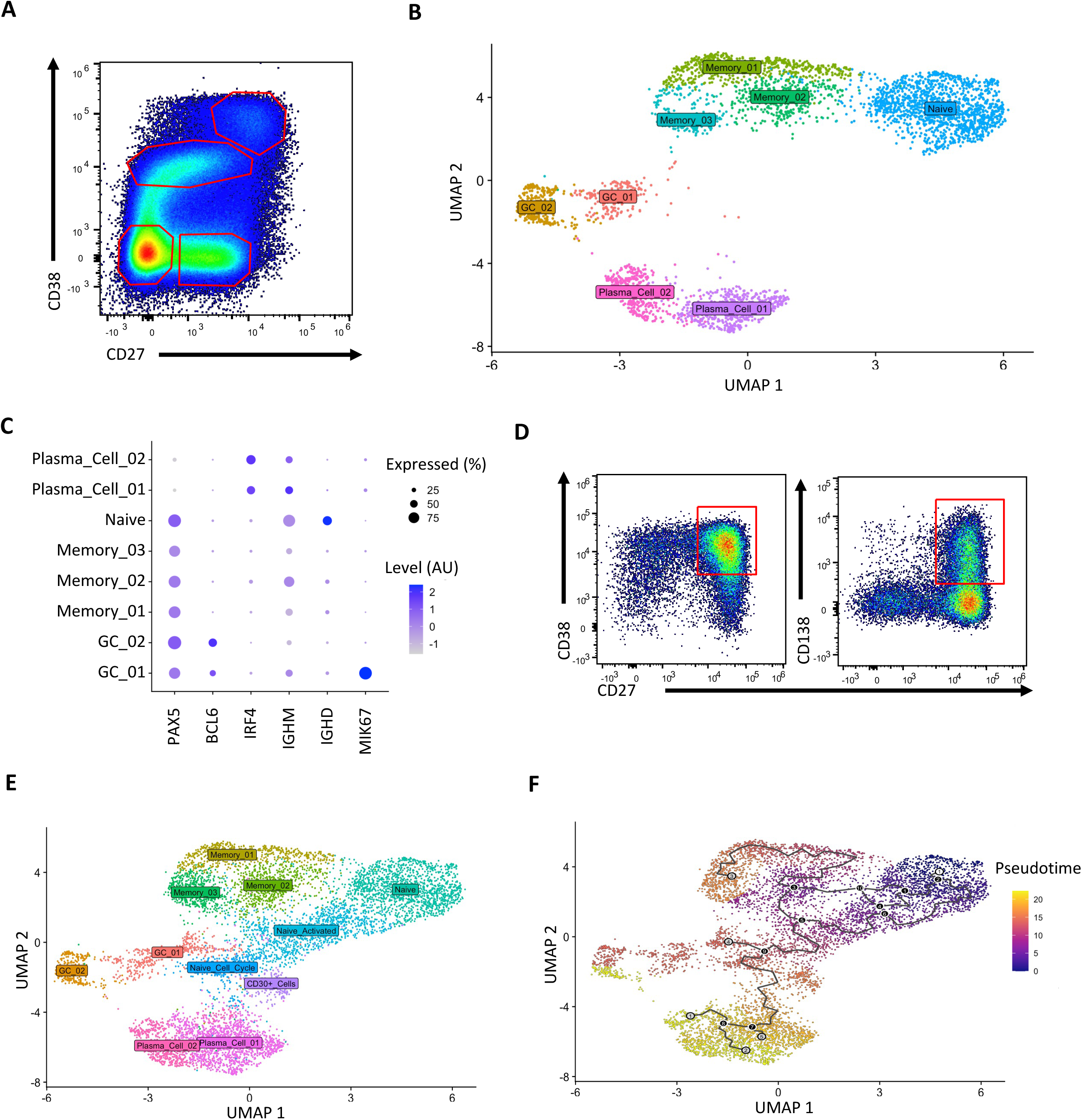
***In vitro* differentiation of primary B cells identifies transitionary populations prior to plasma cell terminal differentiation. (A)** Representative plot of the B cell populations in primary tonsil samples with red gates highlighting Naïve B cells (CD27-CD38-), Memory B cells (CD27+CD38-), Germinal Center B cells (CD27^intermeidate^CD38^intermediate^), and plasma cells (CD27^high^CD38^high^). **(B, C)** Primary CD19+ tonsil B cells (female) were subjected to single cell paired ATAC and RNA sequencing. Cells were embedded in weighted nearest neighbor UMAP space generated using both ATAC and RNA datasets **(B).** Cell identities were assigned based on the expression of known lineage-defining genes and cell cycle genes **(C)**. **(D)** Primary tonsil naïve B cells (female, same donor as in **B** and **C**) were cultured with cytokines and mitogens to induce plasma cell differentiation which was confirmed by staining for CD27+CD38+CD138+ cells. Cells harvested over the course of the differentiation were subjected to the same sequencing as in **(B)** and used to create a new UMAP embedding **(E)** containing both primary and *in vitro* differentiating cells. The integrated primary and *in vitro* datasets were used for pseudotime trajectory analysis with subsequent mapping from the naïve B cell cluster to the plasma cell clusters shown in **(F)**.

Published *in vitro* plasma cell differentiation protocols rely on a combination of IL-4 and lipopolysaccharide (LPS), or combinations of IL-15 or IL-21 with mitogenic stimuli to facilitate differentiation to an antibody secreting state (Hipp et al., 2017; Jourdan et al., 2009; Matsuda et al., 2015; Wang et al., 2018). However, human B cells differ significantly from mouse B cells in the expression of various receptors, including the LPS sensor Toll-Like Receptor 4 (TLR4) (Bekeredjian-Ding and Jego, 2009; Hwang et al., 2009). As such, we chose to combine aspects of these protocols to mimic what might be experienced upon antigen-activation of human B cells. Starting with naïve B cells, the differentiation scheme as depicted in (Sup Fig 1) successfully produced plasma cells by Day 13 of culture as marked by their CD27^+^CD38^+^CD138^+^ surface phenotype (Fig 1D). We harvested cells during *in vitro* differentiation at days 4, 7, 10, and 13 and performed multiome scRNA- and scATAC-seq. Integrating the combined epigenetic and transcriptional profiles of these *in vitro* activated B cells with those of the primary tonsillar B cells revealed many cells populating the intervening regions between naïve B cells and memory B cells, germinal center B cells, and plasma cells (Fig 1E). Using Monocle 3 (Cao et al., 2019), pseudotime analysis was performed to predict changes in cell state and infer developmental trajectories based on gene expression profiles. This analysis mapped from the naïve B cell cluster predominantly to plasma cell cluster 1 while bypassing the germinal center clusters (Fig 1F).

### Human plasma cells exist as CD44v9^+^ and CD44v9^-^ subsets in both tonsil and bone marrow

We next sought to define the differences between the two distinct plasma cell clusters (Fig 1B). These clusters were found to differ in a small number of genes with major differences in expression of *RGS13* and the surface markers CD10 (*MME*), CD31 (*PECAM1*), *CD38*, and *CD44*, which directly modulates plasma cell differentiation *in vivo* (Calderón et al., 2026) (Fig 2A). We next checked the expression of the corresponding proteins flow cytometrically. For CD44, we also specifically assessed the variant 9 splice isoform (CD44v9) given its known role in promoting IL-6-dependent survival in myeloma cells (Van Driel et al., 2002). Consistent with the multiome analysis, we observed a CD31^high^CD38^low^CD44v9^+^ subset corresponding to plasma cell cluster 1, and a CD31^low^CD38^high^CD44v9^-^ subset corresponding to plasma cell cluster 2 in primary tonsil samples (Fig 2B). We also observed bone marrow plasma cell subsets, gated as described in Sup. Fig 2A, that expressed or lacked CD44v9 expression (Fig 2C). We did not observe correlations between CD44v9 and CD19 expression, the latter of which has been suggested to mark either shorter-lived or GC-independent plasma cells (Brynjolfsson et al., 2017; Ferreira-Gomes et al., 2024; Halliley et al., 2015; Landsverk et al., 2017; Mei et al., 2015). Expression of CD44v9 seemed largely confined to the plasma cell subset that also lacked CD10 expression, with only minor staining observed in memory B cells or germinal center B cells (Sup Fig 2B). We observed an increase in IL-6 receptor expression in the CD44v9^+^ tonsil subset (Fig 2D). The CD44v9 isoform specifically is able to induce IL-6 secretion from stromal cells (Van Driel et al., 2002). Given the fact that IL-6 is a known survival and differentiation modulating factor for plasma cells (Jourdan et al., 2014), this suggests a potential functional difference between these subsets which are known to take up residence among IL-6 secreting stromal cells (Wols et al., 2002).

**Fig. 2.**
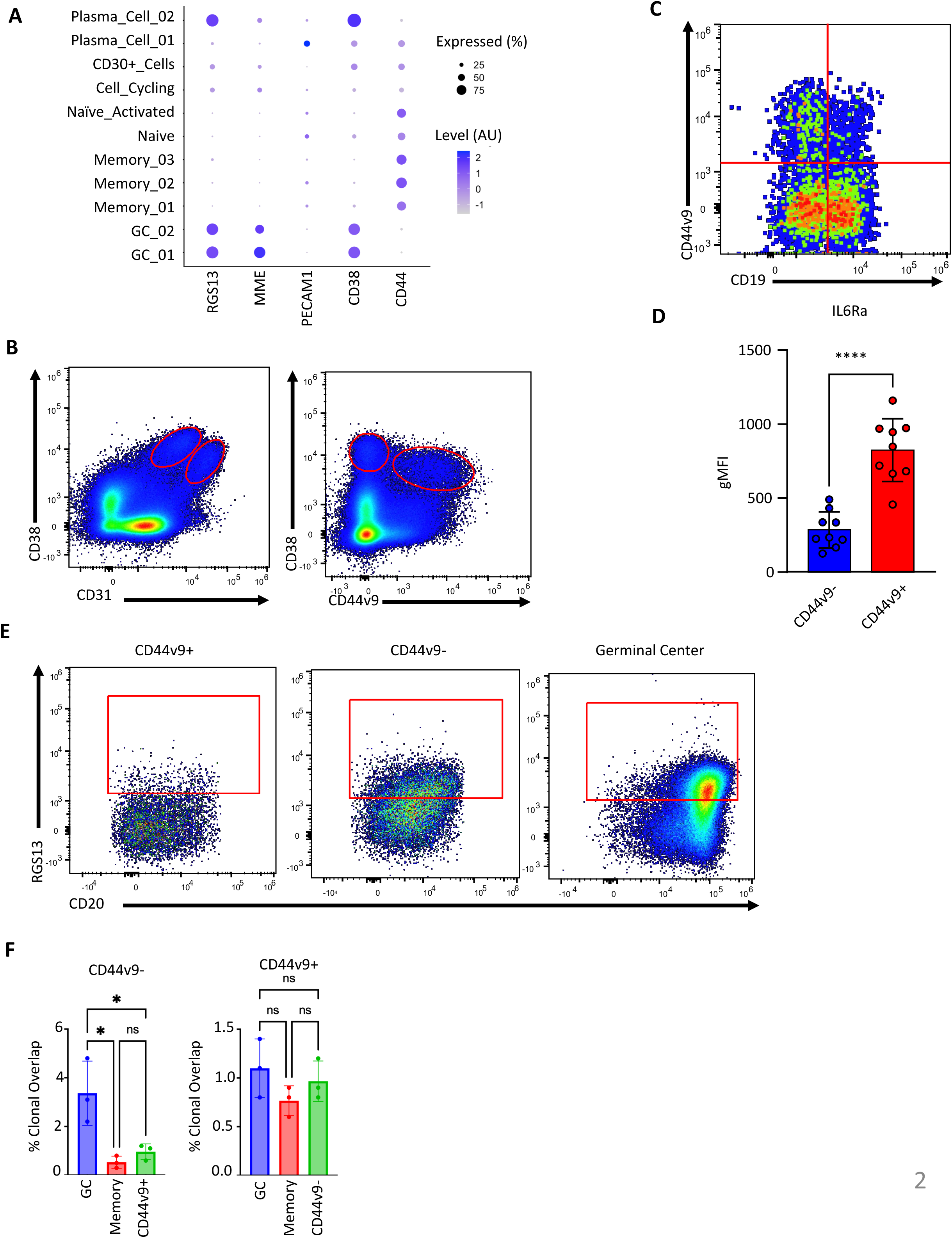
Tonsil plasma cells are comprised of at least two distinct populations. **(A)** Examples of differentially expressed genes between tonsil plasma cell subsets used for flow cytometry validation. **(B)** Representative gating strategy for the two identified plasma cell clusters based on CD31, CD38, and CD44v9. **(C)** CD44v9 protein expression observed in CD27+CD38+ bone marrow plasma cells. **(D)** Differences in IL-6 receptor expression between plasma cell subsets (n = 8 tonsil donors). **(E)** Representative RGS13 protein expression in tonsil plasma cell subsets and germinal center B cells. **(F)** IgH BCR clonal overlap between CD44v9-based tonsil plasma cell subsets, memory B cells, and germinal center B cells (n = 3 tonsil donors). Statistical significance was evaluated using a t-test in figures where only two groups are being compared. In all other instances, one-way analysis of variance (ANOVA) was performed to test for statistical significance. Data are presented as mean ± standard error of the mean (SEM). Statistical significance was defined as *p < 0.05, **p < 0.01, ***p < 0.001, and ****p < 0.0001.

We next confirmed the prediction from our transcriptomic data that the tonsil CD44v9^-^ tonsil plasma cell population expresses higher levels of RGS13, a transcription factor and regulator of G-protein signaling (Landsverk et al., 2017; Xie et al., 2008), similar to that found in germinal center B cells (Fig 2E). RGS13 and CD10 expression suggest that this plasma cell subset may arise through a germinal center origin. Through sorting of specific populations and IgH sequencing, we observed more clonal overlap between CD44v9^-^ plasma cells and germinal center B cells than with memory B cells or CD44v9^+^ plasma cells (Fig 2F). These data are perhaps suggestive of a germinal center origin of CD44v9^-^ plasma cells. However, the overall level of overlap across populations was low, especially when comparing CD44v9^+^ plasma cells against other B cell subsets (Fig 2F), thereby limiting the conclusions that could be drawn. Given that the donor tonsils were all at indeterminant stages of responding to unknown infection(s), we reasoned that focused *in vitro* differentiations may provide more information on the ontogenetic origins, cellular routes, and transcriptional programs that give rise to these plasma cell subsets.

### Human CD44v9^+^ plasma cells can be formed *in vitro* through a transient CD30^+^ intermediate

Pseudotime trajectory analysis of our *in vitro* differentiations did not map to germinal center B cells (Fig 1F). Thus, this system potentially provides a way to define the cellular intermediates and terminal plasma cell fates of a germinal center-independent ontogenetic route. We identified CD30, encoded by the gene *TNFRSF8*, as uniquely expressed in a population of cells inferred to immediately precede plasma cell cluster 1. To validate this finding, protein expression of CD30 along with other known marker genes was assessed by flow cytometry across multiple days of *in vitro* differentiation. Dividing cells into populations based on expression of CD27 and CD30 as depicted in (Fig 3A) revealed that CD30^+^CD27^-^ cells emerge early in the differentiation followed by a wave of CD30^+^CD27^+^ cells, before CD30 expression is lost (Fig 3B). As CD27 is primarily expressed in plasma cells, we took this as evidence of an early transient population of CD30^+^ pre-plasma cells that loses CD30 expression upon further maturation. To confirm this, at Day 4 of culture, we sorted 20,000 *in vitro* differentiating CD30^+^ or CD30^-^ cells to allow them to continue differentiating separately (Fig 3C), and quantified the number of plasma cells they generated (Fig 3D). CD30^+^ cells generated significantly more plasma cells than did CD30^-^ cells. Following a similar approach, we determined that a second developmental stage likely occurs subsequently in which cells that downregulate CD20 expression represent a later-stage precursor population to terminally differentiated plasma cells (Fig 3E-F). Consistent with this interpretation, we observed high levels of IRF4, the master regulator of plasma cell differentiation, in CD20^low^ cells (Sup. Fig. 3A).

**Figure 3:**
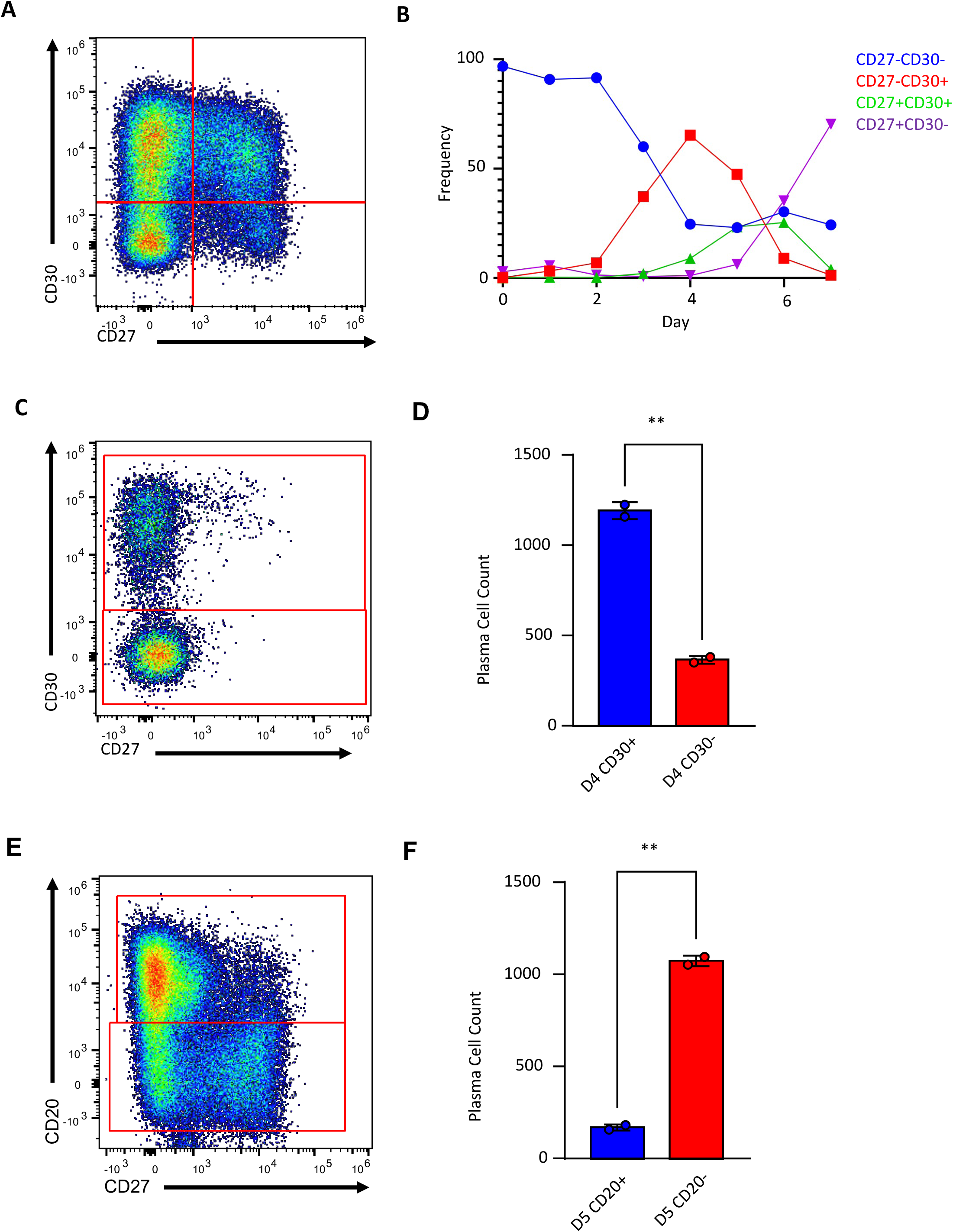
Plasma cell differentiation proceeds through a transient CD30+ intermediate. **(A)** Representative gating scheme for *in vitro* differentiating cells based on CD27 and CD30 expression. **(B)** Quantification of cell populations overtime in culture. **(C)** Representative gating scheme for sorting CD30+ and CD30- populations. **(D)** Plasma cell yields from day 4 sorted CD30+ or CD30- populations (20,000 each, n=2). **(E)** Representative gating for sorting CD20+ and CD20- populations. **(F)** Plasma cell yields from day 5 sorted CD20+ and CD20- populations (20,000 each, n=2). Statistical significance was evaluated using a t-test data are presented as mean ± standard error of the mean (SEM). Statistical significance was defined as *p < 0.05 and **p < 0.01.

Given that the *in vitro* culture system seems to represent a germinal center-independent route to plasma cell differentiation, we sought to test our hypothesis that these cultures would predominantly give rise to CD44v9^+^CD31^high^CD38^low^ plasma cells. Day 4 cultures were investigated as they were expected to have a large number of cells at various intermediate stages of differentiation. By examining patterns of CD27, CD31, CD38, and CD44v9 co-expression, we identified predominant expression of CD31 with CD44v9 while CD27 and CD38 expression was mostly limited to cells that had become CD31^+^CD44v9^+^ (Fig 4A). These data suggest that CD31 and CD44v9 expression temporally precedes CD27 and CD38 expression.

**Figure 4:**
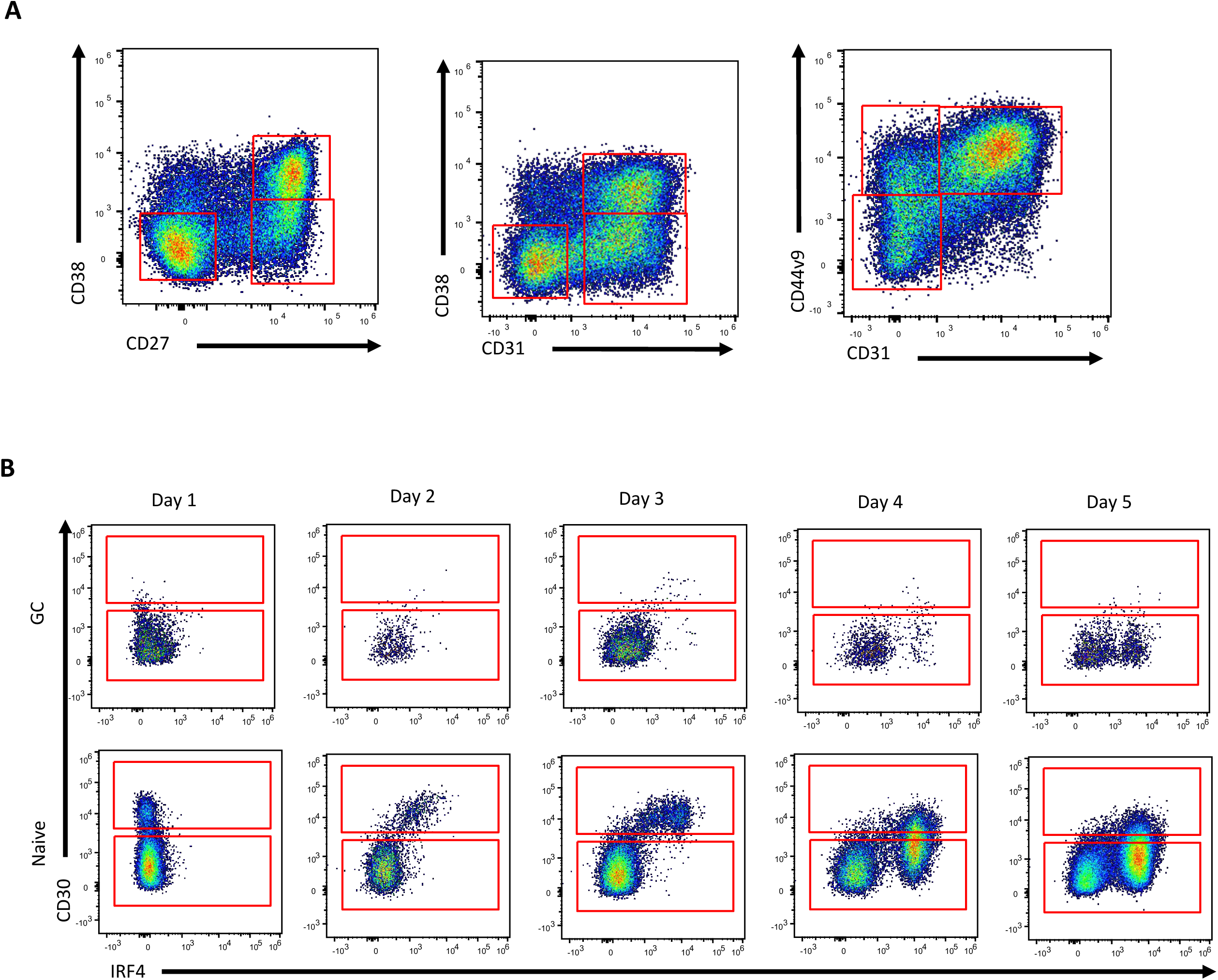
Orderly progression of surface markers during *in vitro* plasma cell differentiation. **(A)** Gating scheme used to define CD27, CD31, and CD38 populations during *in vitro* differentiation of primary naïve B cells. **(B)** Comparison of CD30 expression between *in vitro* differentiating germinal center or naïve B cells and its correlation with IRF4 expression.

Next, we sought to characterize germinal center cell-dependent plasma cell differentiation in this system, with the prediction that this route would predominantly yield CD44v9^-^CD38^high^ cells and might not traverse through a CD30^+^ stage. Starting from sorted primary germinal center B cells, we found no evidence of a CD30^+^ intermediate (Fig 4B). The small fraction of germinal center B cells that reached high IRF4 expression lacked CD30 expression, while CD38 expression (normally expressed by germinal center B cells) was maintained early on (Sup. Fig. 3A-B). Of the plasma cells that were formed, a substantial fraction were CD44v9^-^ (Sup. Fig. 3C-D). Though some germinal center-derived plasma cells were CD44v9^+^, the frequency of these cells was substantially less than those generated from naïve B cells (Sup. Fig. 3C-D). These data suggest the two primary plasma cell subsets we identified may predominantly represent GC-independent and GC-dependent origins.

Memory B cells tend to generate plasma cells mostly, though not exclusively, through a germinal center-independent route *in vivo* (Mesin et al., 2020; Turner et al., 2020; Wong et al., 2020). We therefore repeated these differentiations with purified naïve and memory B cells (CD27^-^ atypical and CD27^+^ canonical) sorted as depicted in (Fig 5A) to determine whether memory B cells progress through a CD30^+^ intermediate. Memory B cells were also able to form CD30^+^ intermediates in greater numbers and more rapidly than did naive B cells, (Fig 5B), consistent with what is observed *in vivo* (Schiepers et al., 2023). To gain insights as to how memory B cells generate plasma cells more efficiently than do naïve B cells, we tested the signals required to generate the CD30^+^ intermediate cells. CD30^+^ cells appeared when TLR9 signaling was paired with CD40L, IL-10, or IL-21 with compounding effects when combined further (Fig 5C). Flow cytometry analysis revealed that memory B cells express higher levels of the IL-10 receptor and TLR9 than do naïve B cells (Fig 5D), potentially underlying the heightened responsiveness of memory B cells.

**Figure 5.**
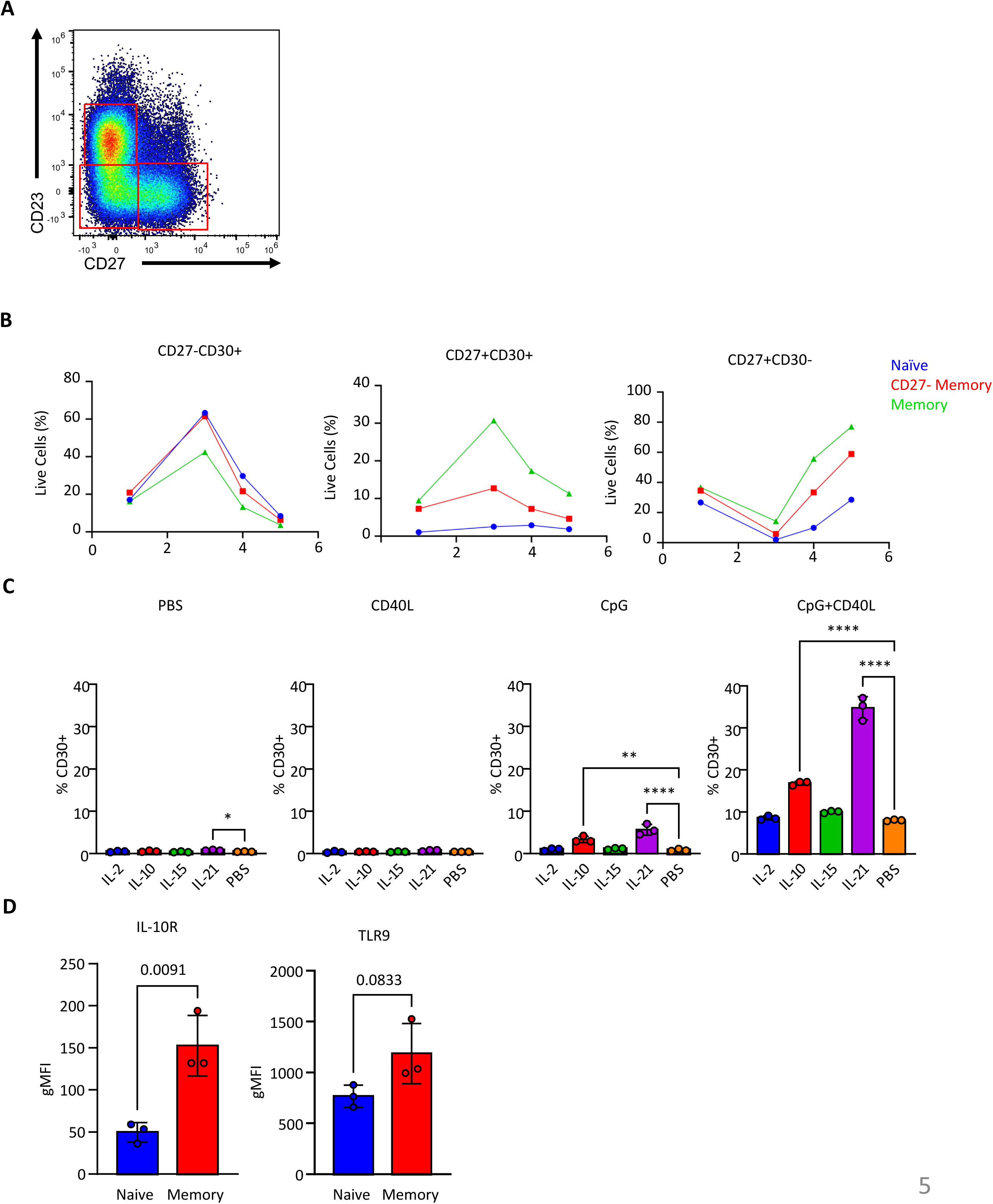
Identification of formation requirements for CD30+ cells and comparison of naïve and memory B cell differentiation. **(A)** Representative gating scheme for sorted naïve (CD23+CD27-), atypical (CD23-CD27-), and canonical (CD23-CD27+) memory B cells used in **(B)**. **(B)** Changes in CD27 and CD30 populations over time for *in vitro* differentiating naïve, atypical, or canonical memory B cells (20,000 seeded each, n = 1). **(C)** Formation of CD30+ cells in response to individual cytokines and mitogens or their combinations (n = 3 separately seeded wells, all from 1 tonsil donor). **(D)** Comparison of IL-10R and TLR9 between naïve and memory B cells (n = 3 tonsil donors). Statistical significance was evaluated using a t-test in figures where only two groups are being compared. In all other instances, one-way analysis of variance (ANOVA) was performed to test for statistical significance. Data are presented as mean ± standard error of the mean (SEM). Statistical significance was defined as *p < 0.05, **p < 0.01, ***p < 0.001, and ****p < 0.0001.

### Cellular intermediates corresponding to *in vitro* cultures exist *in vivo*

Armed with the detailed characterization of how markers change in our *in vitro* system over time, we sought to validate the existence of the corresponding populations in primary tonsils. To circumvent the issues posed by the rarity of CD30^+^ cells in primary samples, five individual tonsil donors were used to analyze approximately 10 million total B cells each. As our *in vitro* system predicts CD30 to be expressed first, followed by expression of CD44v9 before CD30 is eventually turned off, we gated the tonsil B cells into CD30^+^CD44v9^-^, CD30^+^CD44v9^+^, and CD30^-^CD44v9^+^ populations with a representative plot and fluorescence minus one (FMO) controls shown in (Fig 6A). As observed for our *in vitro* system, the CD30^+^CD44v9^-^ population was predominately CD20^+^ with no CD31 or CD38 expression (Fig 6B). We observed CD27 expression in the CD30^+^CD44v9^+^ population, similar to our *in vitro* system (Fig 6B). The predicted most mature CD30^-^CD44v9^+^ cells continued this trend, with CD20 being largely absent from this population and having high expression of CD31 and CD38 (Fig 6B). Crucially, this recapitulated the finding from our *in vitro* system where high CD38 expression only occurs in the cells that have differentiated to the CD30^-^ stage where they start to resemble fully mature plasma cells. We additionally confirmed that the expression of CD44v9 is partially linked to CD30 *in vitro*, exhibiting a correlated expression until the cells downregulate CD30 while maintaining CD44v9 as plasma cells (Sup. Fig. 5). Together, these data confirm that the populations we identified from our *in vitro* cultures correspond to primary cellular intermediates *en route* to plasma cells *in vivo*.

**Figure 6.**
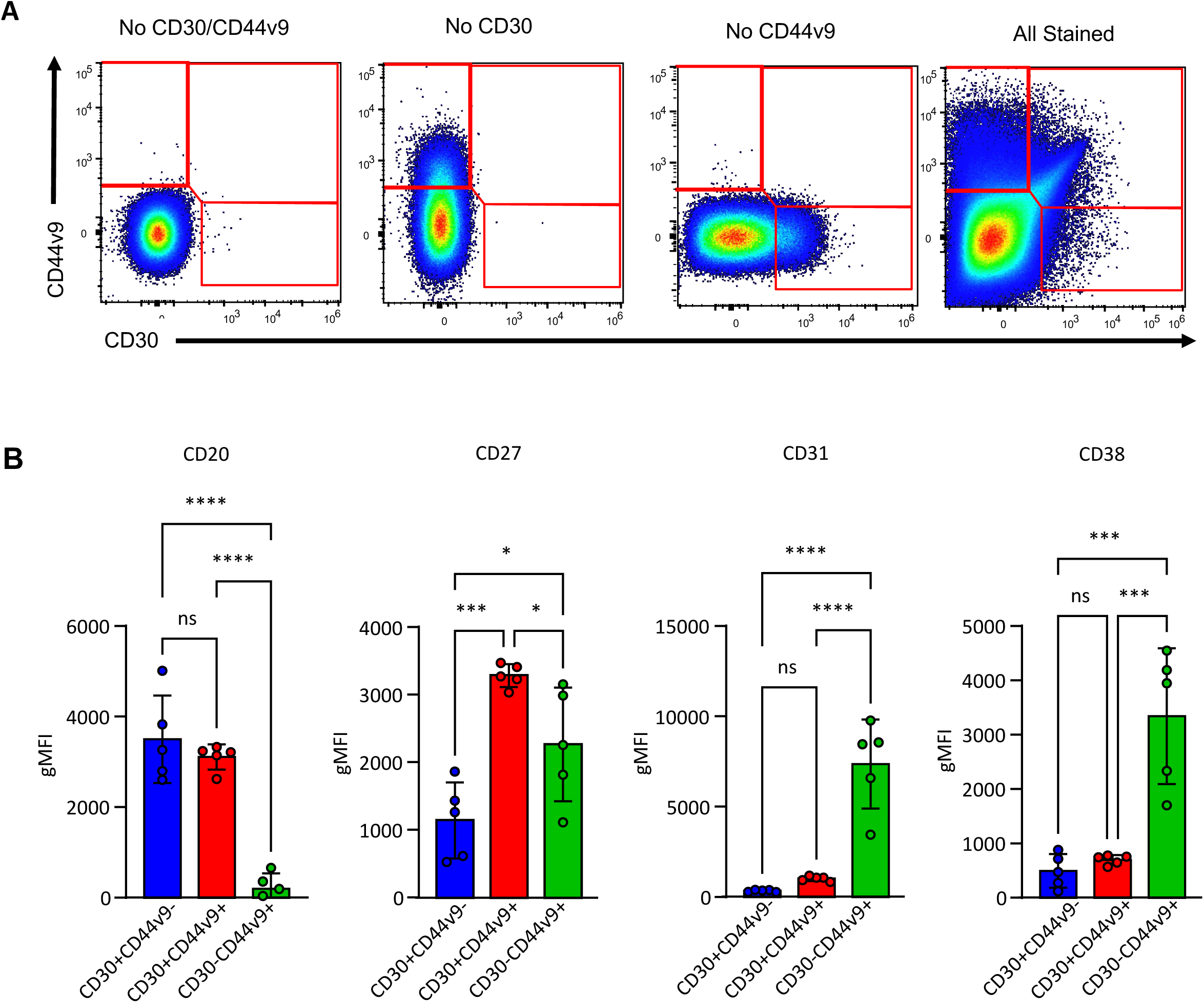
Comparison of primary CD30 and CD44v9 populations. **(A)** Representative gating scheme for comparing CD30 and CD44v9 populations in **(B)** highlighting the lack of compensation artefacts with fluorescence minus one and minus two controls. **(B)** Comparison of CD20, CD37, CD31, and CD38 expression between the subsets identified in **(A)** (n = 5 tonsil donors). Statistical significance was evaluated by one-way analysis of variance (ANOVA). Data are presented as mean ± standard error of the mean (SEM). Statistical significance was defined as *p < 0.05, **p < 0.01, ***p < 0.001, and ****p < 0.0001.

### Gene regulatory network inference identifies actionable transcriptional regulators of plasma cell differentiation

Having characterized the cellular intermediates leading to distinct plasma cell fates, we sought to identify transcriptional regulators that could be pharmacologically targeted to enhance differentiation efficiency. The transient nature of the CD30+ intermediate *in vivo* suggested tight transcriptional control, presenting an opportunity to identify rate-limiting factors. Though the relative inefficiency of our *in vitro* system may have allowed us to observe and characterize these cellular intermediates, we reasoned that improving its efficiency may better align the signals and programs that occur *in vivo.* Starting with this *in vitro* system, we therefore sought to identify regulators of plasma cell differentiation. We employed CellOracle (Kamimoto et al., 2023), to construct gene regulatory networks (GRNs) from our integrated scRNA-seq and scATAC-seq data. This approach identifies transcription factors (TFs) based on three complementary metrics: degree centrality (number of regulatory connections), betweenness centrality (control of information flow through the network), and eigenvector centrality (influence through connections to highly-connected nodes). We focused on betweenness centrality to identify TFs that serve as critical regulatory gatekeepers controlling transitions between cell states. We compared the predicted transcriptional regulators of the CD30^+^ cell cluster and the plasma cell 1 cluster depicted in (Fig 1E). This comparison (Fig 7A) identified enhanced activity of known plasma cell lineage-defining transcription factors IRF4, Blimp-1, and XBP1, and predicted that the CD30^+^ identity could be regulated by STAT1, POU2F2, and MEF2C. Bioinformatics simulations for overexpression and knockout of STAT1, XBP1, IRF4, and MEF2C are depicted in Sup. Fig 6.

**Figure 7:**
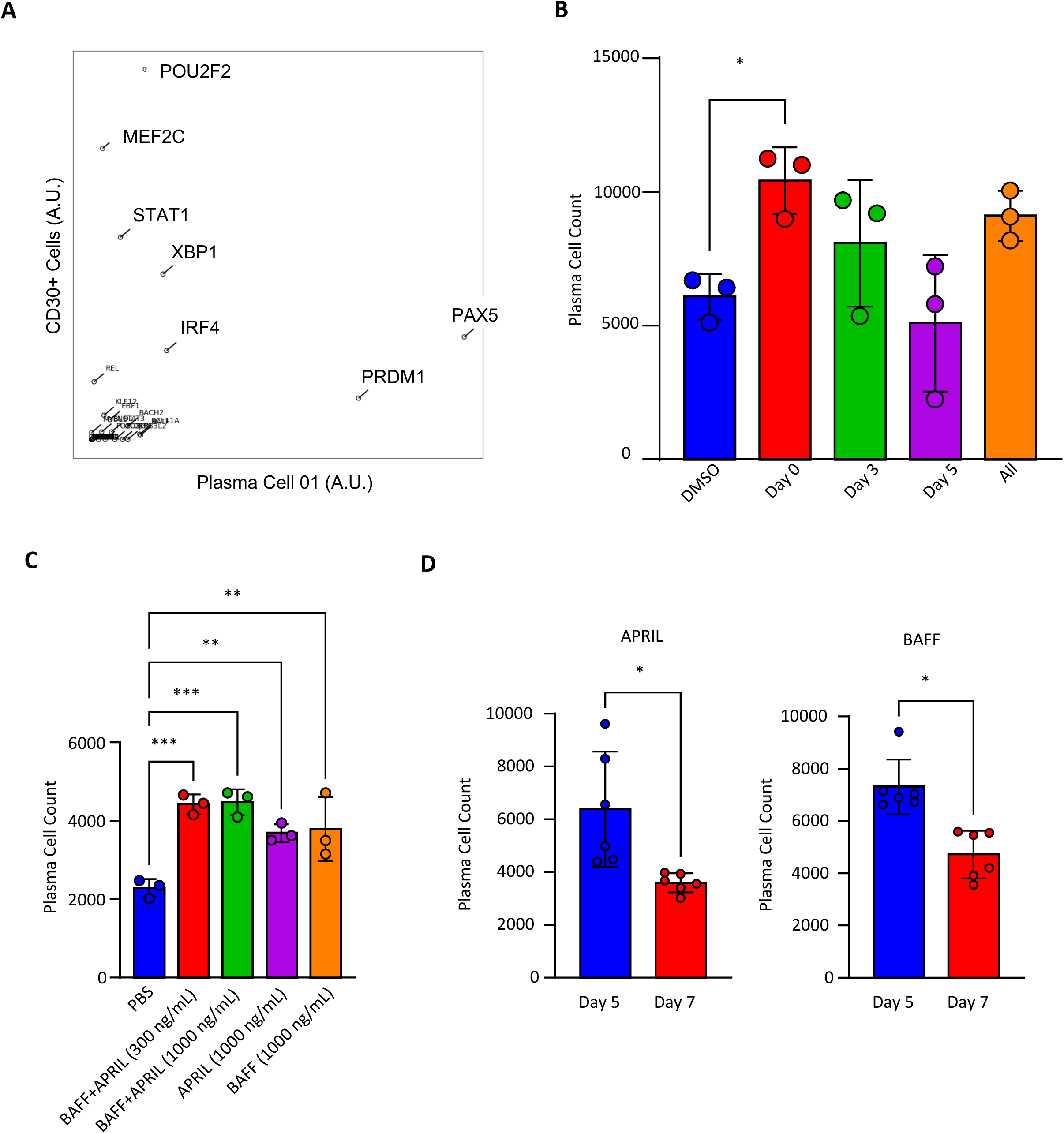
MEF2C, BAFF, and APRIL act prior to terminal plasma cell differentiation. **(A)** Betweenness centrality for transcriptional regulators determined by CellOracle analysis of the CD30+ Cells and Plasma Cell 01 clusters depicted in Fig 1F. Final plasma cell yields for **(B)** A366 added on Day 0, 3, 5, or each day (n = 3), **(C)** BAFF and APRIL given alone or in combination on Day 5 (n = 3), and **(D)** BAFF and APRIL given alone on Day 5 or Day 7 (n = 6). Statistical significance was evaluated using a t-test in figures where only two groups are being compared. In all other instances, one-way analysis of variance (ANOVA) was performed to test for statistical significance. Data are presented as mean ± standard error of the mean (SEM). Statistical significance was defined as *p < 0.05, **p < 0.01, and ***p < 0.001.

To functionally validate our GRN predictions, we prioritized MEF2C due to conflicting reports on its role in B cell responses. Previous studies showed that genetic deletion of MEF2C in mouse B cells impaired early development (Debnath et al., 2013) but did not affect CD40L and TLR9 responses (Wilker et al., 2008), which are the exact signals we identified as critical for CD30+ intermediate formation. This apparent contradiction suggested context-dependent functions that warranted investigation. MEF2C activity can be pharmacologically enhanced by inhibiting the lysine methyltransferase G9a, which normally suppresses MEF2C function (Ow et al., 2016). Treatment with the small molecule G9a inhibitor A366 significantly increased plasma cell yields when provided early in differentiation (Day 0-3), whereas later addition had no effect (Fig 7B). This temporal specificity aligns with the transient appearance of the CD30+ intermediate and suggests MEF2C activity is required during the initial commitment phase.

To confirm that A366 effects were mediated through MEF2C, we performed scRNA- and ATAC-seq on treated versus control cells. While global transcriptional changes were modest, the differentially expressed genes primarily involved mitochondrial respiration (Sup. Fig 7A). Importantly, these changes overlapped significantly with genes reported to be dysregulated in MEF2C knockout cells (Sup. Fig 7B) (Ow et al., 2016), confirming on-target effects. These findings establish MEF2C as a novel, pharmacologically accessible regulator of human plasma cell differentiation.

### Coordinated activity of multiple transcription factors drives plasma cell maturation

Having established the role of MEF2C in the CD30+ intermediate, we functionally validated additional predicted regulators to define the complete transcriptional program. XBP1, activated in response to ER stress from antibody secretion (Gaudette et al., 2020), could be engaged earlier using the PPARγ agonist Rosiglitazone, which activates the integrated stress response (Garcia-Bates et al., 2009). This treatment increased plasma cell yields when added at Day 3 (Sup. Fig 8A), coinciding with the onset of antibody secretion.

IRF4, a master regulator of plasma cell differentiation, is normally regulated through degradation by the E3 ubiquitin ligase Cbl-b (Li et al., 2018). Treatment with the Cbl-b inhibitor NX-1607 (Gallotta et al., 2022) at Day 5 enhanced plasma cell yields (Sup. Fig 8B), confirming continued importance of IRF4 during later-stage differentiation. Finally, STAT1 activation through interferon-γ (Zumaquero et al., 2019) combined with the STAT1-enhancing small molecule 2-NP (Lynch et al., 2007) also increased plasma cell yields when provided at Day 5 (Sup. Fig 8C).

Notably, each factor showed distinct temporal requirements, revealing a coordinated transcriptional program: MEF2C acts early during CD30+ intermediate formation (Days 0-3), XBP1 engages at the onset of antibody secretion (Day 3), and IRF4/STAT1 continue to drive terminal differentiation (Day 5+). This sequential activation pattern suggests a multi-stage regulatory cascade rather than simultaneous engagement of all factors.

### Temporal coordination of signals defines an optimized differentiation protocol

Beyond transcription factor regulation, we investigated survival and differentiation signals. Despite reported roles in plasma cell survival (Jourdan et al., 2014), BAFF and APRIL showed their primary effects when added at Day 5 during the late pre-plasma cell stage rather than after terminal differentiation (Fig 7C-D). This unexpected finding suggests these factors may function primarily as differentiation signals in human plasma cells rather than pure survival factors for mature cells.

Integrating our findings on temporal requirements for transcription factors, cytokines, and survival signals, we constructed a comprehensive model for human plasma cell differentiation through the GC-independent, CD30+ intermediate route (Sup. Fig 9A). This model reveals three distinct phases: (1) initial activation and CD30+ intermediate formation (Days 0-4) requiring TLR9, CD40, IL-10/IL-21 signals and MEF2C activity; (2) antibody secretion initiation (Days 3-5) requiring XBP1 engagement and BAFF/APRIL signaling; and (3) terminal differentiation (Days 5+) driven by sustained IRF4 and STAT1 activity. We also provide a detailed surface marker expression timeline (Sup. Fig 9B) to guide identification of differentiation intermediates.

Importantly, this temporal roadmap provides actionable strategies for enhancing plasma cell generation in vitro, with direct applications to therapeutic antibody production and vaccine development.

## Discussion

A central goal in immunology is to control B cell differentiation with sufficient precision to enhance vaccine responses or generate therapeutic antibodies. Our identification of MEF2C, STAT1, and POU2F2 as regulators of a critical CD30+ intermediate, combined with validation that their pharmacological modulation enhances plasma cell yields, represents a significant advance toward this goal. Notably, MEF2C had not previously been implicated in this pathway despite extensive study of plasma cell differentiation, highlighting the power of integrating multiomic profiling with functional validation. The sequential temporal requirements we defined for MEF2C (early), XBP1 (intermediate), and IRF4/STAT1 (late) provide a roadmap for optimizing in vitro plasma cell generation, which is important relevant to both basic research and cellular engineering applications. The broader significance lies in demonstrating that computational inference from multiomic data, when rigorously validated, can identify non-obvious therapeutic targets. While IRF4, XBP1, and PRDM1/Blimp-1 were already known as plasma cell regulators, the role of MEF2C emerged specifically from our GRN analysis and would not have been predicted from prior literature. This underscores the value of unbiased network approaches in discovering context-dependent regulatory mechanisms

Irrespective of their ontogeny and activation cues that led to their formation, plasma cells can be readily identified by common transcriptional programs. Yet there is considerable heterogeneity within the plasma cell pool on the basis of longevity (Amanna et al., 2007; White et al., 2015), isotype (Higgins et al., 2022; Tellier et al., 2024b; Vecchione et al., 2024), tissue of formation or residence (DiLillo et al., 2008; Ise et al., 2025; Kim et al., 2013; Salmi and Jalkanen, 2005; Tellier et al., 2024b; van Spriel et al., 2012), and functional properties (Fritz et al., 2012; Lam et al., 2018; Tellier et al., 2024b), with much likely left to be discovered. While very useful biologically, this richness complicates our ability to unravel the mechanisms that give rise to the diverse pool of plasma cells critical for mediating humoral immunity. Inroads have been made into the ontogeny of mouse plasma cells through new sequencing technologies and lineage tracing and genetic manipulation (Gómez-Escolar et al., 2022; Schulz et al., 2025). However, the limitations of working with human samples leave open many questions including whether the same regulatory mechanisms and cellular routes of differentiation described for mice hold true. Using single cell multiomic sequencing of both primary and *in vitro* differentiating human plasma cells and functional validation, we have identified multiple routes B cells can take on their way to at least two distinct plasma cell fates.

The GC-independent route we have described here, proceeding through a CD30^+^ intermediate, gives rise to predominately CD44v9^+^ plasma cells while GC-dependent routes appear biased to CD44v9^-^ plasma cells. However, it remains an open question if this is truly a clear cut distinction rather than simply a biasing towards one fate, similar to how M1 and M2 polarized macrophages are now recognized as extreme ends of a more continuous spectrum observed *in vivo* (Mosser and Edwards, 2008; Murray et al., 2014; Orecchioni et al., 2019). As the extrinsic factors we identified as required for proceeding through the CD30^+^ route are expected to also be present within or around germinal centers, there is considerable potential for intermingling between these two possible routes with unknown additional layers of regulation likely at play *in vivo*.

While the exact path a B cell takes to terminal plasma cell remains unclear, what is more certain is the potential for functional differences between the eventual identity it takes. Adding to previously described differences in longevity and functionality between plasma cell subsets, the subsets we identify here differ in their ability to respond to a known differentiation modulating and survival factor, IL-6 (Jourdan et al., 2014). This difference is even more striking as the basis for identifying them, CD44v9 expression, is known to directly modulate the secretion of IL-6 by stromal cells (Van Driel et al., 2002). Additionally, while CD19 expression has been recently shown to be a marker of past IL-21 exposure rather than longevity (Ferreira-Gomes et al., 2024), the cells we form in the presence of IL-21 still have surface CD19 protein while largely lacking detectable CD19 RNA in our sequencing datasets, further complicating our understanding of primary cells.

Beyond unraveling details of plasma cell differentiation on the basic science side, there are intriguing applications for the discoveries made in this process. The transcriptional regulatory mechanisms we identified have immediate practical applications. For vaccine development, enhancing plasma cell differentiation could improve antibody responses in immunocompromised populations or enable dose-sparing strategies. For therapeutic antibody production, optimizing differentiation protocols using MEF2C activation and temporally-defined factor combinations could increase yields while maintaining antibody quality. The discovery that BAFF and APRIL function primarily during differentiation rather than mature cell survival may explain variable clinical efficacy of BAFF/APRIL-blocking therapies (Cheekati and Murakhovskaya, 2024; Meng et al., 2025; Tian et al., 2025) and suggests that timing of intervention is critical. CD30^+^ B cells have been implicated in various autoimmune diseases and cancers (Higashioka et al., 2020; Weniger et al., 2018). In these contexts, CD30^+^ B cells may be the direct disease-causing agent or necessary intermediate to promote a disease process, suggesting a potential for CD30-depleting therapies in these settings.

## Materials and Methods

### Isolation of Primary B Cells and Plasma Cells

Tonsil samples were obtained from an individual undergoing elective tonsillectomy (Banner-University Medical Center). Tonsils were extracted from anesthetized individuals using Bovie electrocautery in a standard tonsillectomy surgery. Bilateral tonsils were subsequently combined and provided as a fresh sample for processing to the lab with no identifying information. Naïve B cells were isolated using Naive B Cell Isolation Kit II, human (Miltenyi, cat 130-091-150) according to the manufacturer’s protocol with the following modifications: biotinylated antibodies targeting IgG, IgA, IgE, and CD38 were added during the non-B cell labeling step to facilitate removal of atypical class-switched cells as well as contaminating CD27-CD38+ germinal center cells. Bone marrow samples were obtained from individuals undergoing robotics-assisted hip arthroplasty (Tucson Orthopedic Institute). All donors were anonymous, and no patient data was collected as part of this investigation. This work was approved and deemed non-human subjects research by the University of Arizona Human Research Protections Office.

### Cell Culture and Differentiations

Primary B cells were cultured in IMDM supplemented with 10% FBS and Antibiotic-Antimycotic at 37C with 5% CO2, typically in 96-well U-bottom cell culture plates. Cytokine concentrations used for differentiation were determined through titrations to establish working concentrations and added according to the scheme depicted in Fig 1D except where indicated otherwise in the text with the following working concentrations: a-IgM F(ab’)2 (100 ng/mL, Source), IL-2 (50 ng/mL, Stem Cell Technologies), IL-6 (50 ng/mL, Stem Cell Technologies), IL-10 (50 ng/mL, Stem Cell technologies), IL-15 (100 ng/mL, Stem Cell Technologies), IL-21 (25 ng/mL, Stem Cell Technologies), sCD40L (50 ng/mL, Acro Biosystems), ODN 2006 (CpG) (1000 ng/mL, Stem Cell Technologies), IFNa2b (100 ng/mL, Stem Cell Technologies).

### Flow Cytometry Analysis

Cryopreserved primary cell samples were thawed, washed with PBS, and resuspended as single cell suspensions in PBS supplemented with 5% adult bovine serum (FACS buffer). Extracellular staining was performed with titrated concentrations of antibodies targeting the indicated markers for 15-20 minutes on ice after which the cells were washed and resuspended in FACS buffer. For intracellular staining, cells stained for extracellular markers were fixed with 2% paraformaldehyde (PFA) in PBS for 3 minutes at room temperature. The PFA was neutralized with the addition of an equal volume of FACS buffer. Fixed cells were washed and resuspended in PBS with 0.1% saponin to facilitate permeabilization along with titrated concentrations of antibodies targeting the indicated markers on ice for 15-20 minutes. The cells were then washed with 0.1% saponin and resuspended in FACS buffer for analysis. Cells were analyzed on an Thermo Fisher Attune cytometer. Fluorescence activated cell sorting (FACS) was performed on a 5-laser BD FACS ARIA III.

The following antibody clones were used (all purchased from Biolegend unless indicated otherwise): CD10 (clone HI10a), CD19 (clone HIB19), CD20 (clone 2H7), CD21 (clone Bu32), CD23 (clone EBVCS-5), CD27 (clone O323), CD30 (clone BY88), CD31 (clone WM59), CD38 (clone HIT2), CD44 (clone BJ18), CD44v9 (clone RV3), CD138 (clone MI15), RGS13 (Santa Cruz Biotechnology clone G-7 catalog sc-514590 PE), IRF4 (clone IRF4.3E4), Blimp-1(BD Bioscience clone 6D3). DAPI for viability staining was purchased from Millipore Sigma (catalog D9542).

### Combined single-cell ATAC-seq and RNA-seq

Single-cell suspensions were generated from FACS purified primary tonsil cells, bone marrow plasma cells, or *in vitro* differentiating cells (Days 4, 7, 10, and 13). Primary tonsil cells were pooled to give an equal ratio of naïve B cells, memory B cells, germinal center B cells, and plasma cells. 10,000 nuclei per sample were processed and isolated using the Nuclei EZ Prep kit (Sigma-Aldrich) according to the manufacturer’s protocol. Libraries for both ATAC-seq and RNA-seq were prepared using the Chromium Next GEM Single Cell Multiome ATAC + Gene Expression Reagent Bundle (10x Genomics, PN-1000283). Libraries were prepared with i7 indices, multiplexed, and sequenced in partial lanes of the Illumina NovaSeq X Plus Series (PE150) to obtain 25,000 paired-end reads per nuclei. Unique sequences in each i7 index were then used for demultiplexing. Raw sequencing data from both ATAC-seq and RNA-seq were processed using Cell Ranger 2.0.2 (10x Genomics) for alignment to the hg38 reference genome, filtering, and generating expression and accessibility matrices. These matrix files were then imported into RStudio for data processing which includes filtering out low quality cells, normalizing gene expression data using SCTransform and DNA accessibility data using latent sematic indexing (LSI). We then integrated all datasets into a single merged Seurat object following a previously described pipeline (https://stuartlab.org/signac/articles/pbmc_multiomic). We performed linear dimensional reduction on the integrated dataset to create Uniform Manifold Approximation and Projection (UMAP) graphs to visualize the clustering of cells based on both chromatin accessibility and gene expression profiles. Pseudotime analysis was then conducted using Monocle3 (v1.3.1). Cells were ordered based on their differentiation status from primary naïve B cells through primary plasma cells. Cell trajectories were visualized on the UMAP embedding.

### Gene regulatory network (GRN) and perturbation modeling using CellOracle

CellOracle (v2.3.2) was used to infer transcription factor-driven regulatory networks. CellOracle infers TF/regulator activity from the gene expression data using the CellOracle supplied base GRN. Next, these regions are scanned for known transcription factor (TF) binding motifs, which define all possible TF-target gene interactions. CellOracle then integrates scRNA-seq data to retain only TF-target interactions supported by observed expression patterns. This results in dataset-specific gene regulatory networks (GRNs) that were used to identify TFs essential for development of a particular cell cluster. In silico perturbation simulations, overexpression (OE) and knockout (KO), were then used to predict regulatory impacts of a TF on differentiation trajectories.

### Statistical Analysis

Statistical significance was evaluated using a t-test in figures where only two groups are being compared. In all other instances, one-way analysis of variance (ANOVA) was performed to test for statistical significance. Data were presented as mean ± standard error of the mean (SEM). Statistical significance was defined as *p < 0.05, **p < 0.01, ***p < 0.001, and ****p < 0.0001.

## Data Availability

The datasets generated are deposited on NCBI Gene Expression Omnibus under Series GSE318326. All other relevant data supporting the key findings of this study are available within the article and its supplementary files. Source data are provided with this paper.

## ACKNOWLEDGEMENTS

This work was supported by NIH grants R01AI099108 (D.B.), R01AI129945 (D.B.), R21AI176305-01A1 (A.B.), and P30CA023074 (for the Flow Cytometry Shared Resource). This work was also supported by Gates Foundation grants INV-051356, INV-071091, and INV-007910 (D.B.).

---

**Sup. Fig. 1:**
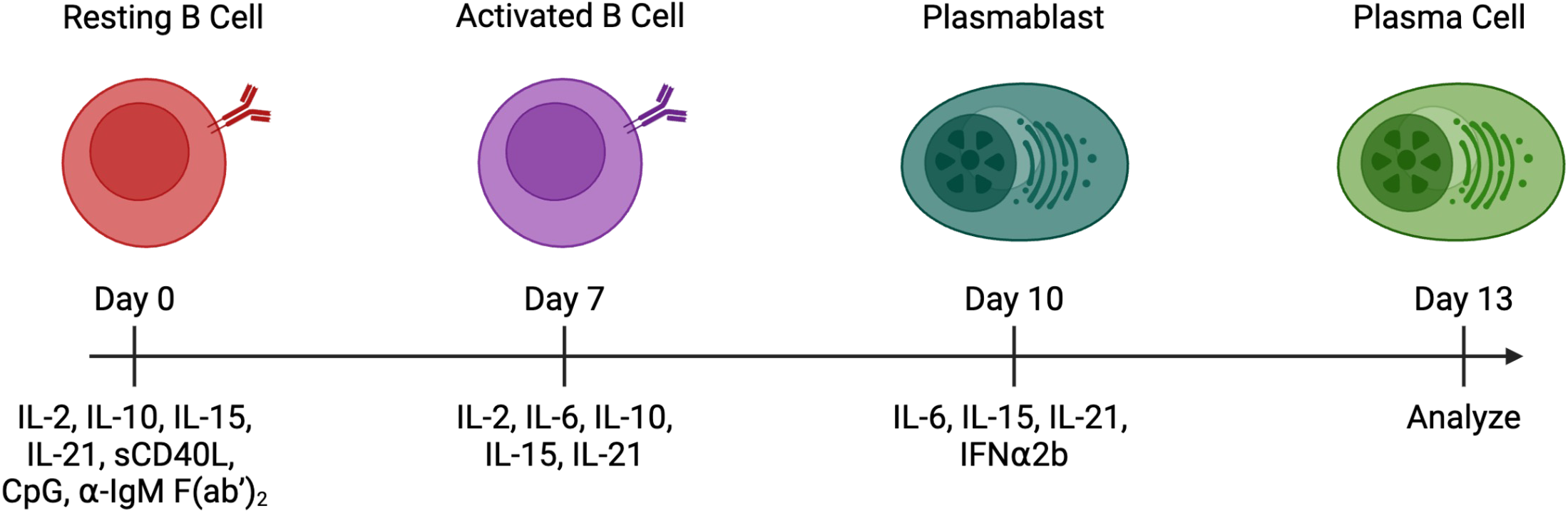
***In vitro* differentiation scheme.** Primary tonsil naïve B cells were cultured with the indicated cytokines and mitogens to induce plasma cell differentiation.

**Sup. Fig. 2.**
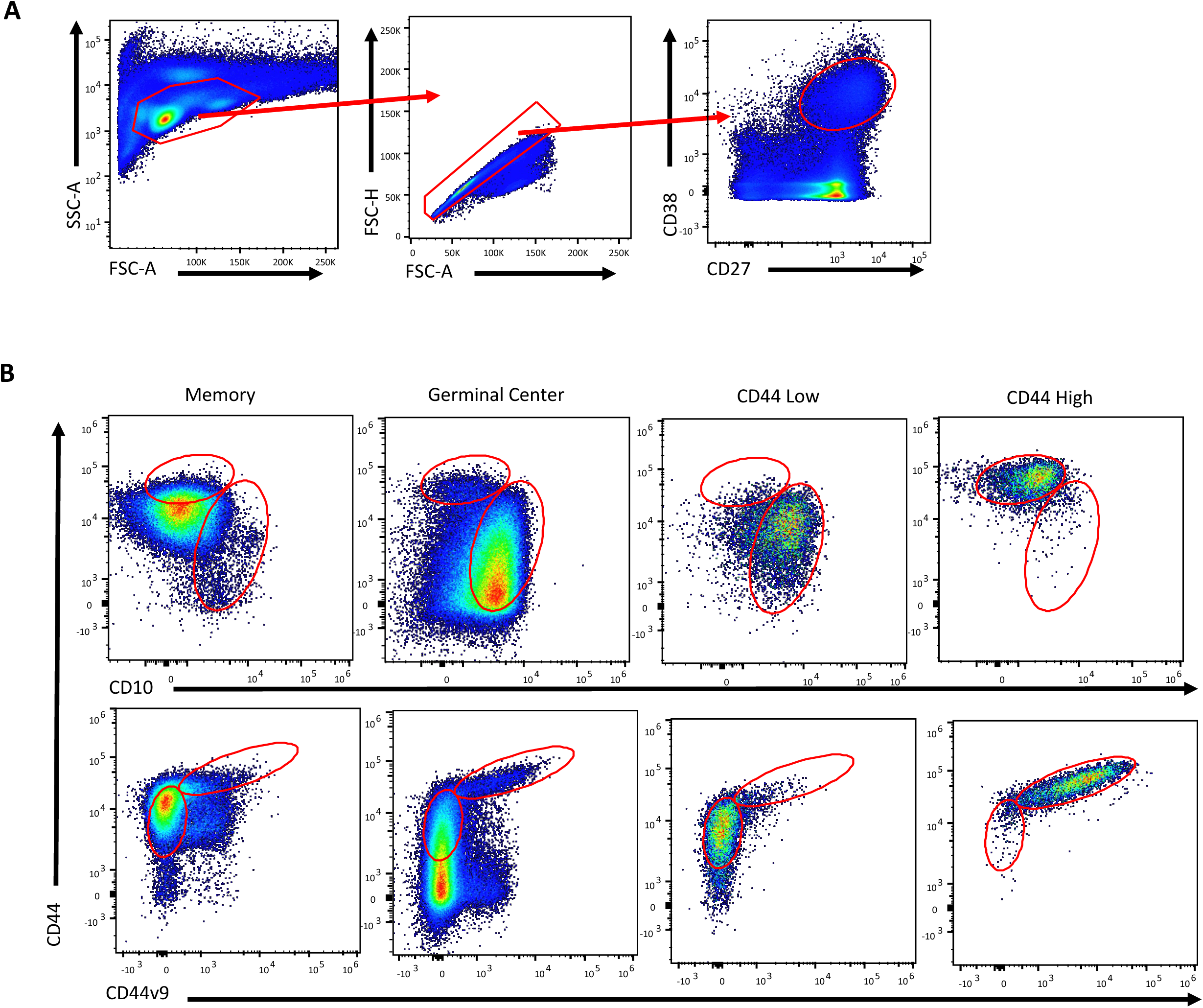
Gating strategies for identification and comparison of tonsil plasma cell subsets. **(A)** Gating scheme for bone marrow plasma cells (CD27+CD38+). **(B)** Comparison of CD44 and CD44v9 expression between tonsil plasma cell subsets and memory and germinal center B cells (n = 1 tonsil donor).

**Sup. Fig. 3:**
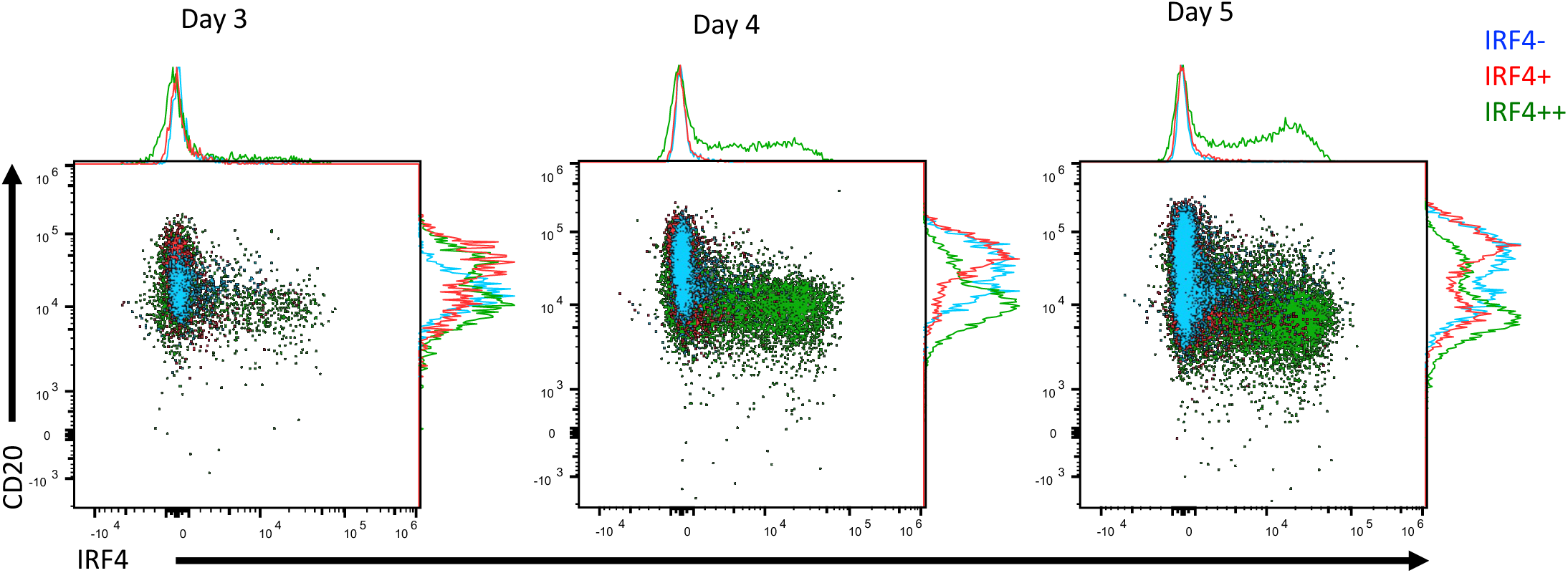
Changes in IRF4 and CD20 observed during *in vitro* differentiation. **(A)** Comparison of CD20 expression in cells with varying levels of IRF4 expression during Day 3, 5, and 5 of *in vitro* differentiation staring from primary naïve B cells.

**Sup. Fig. 4:**
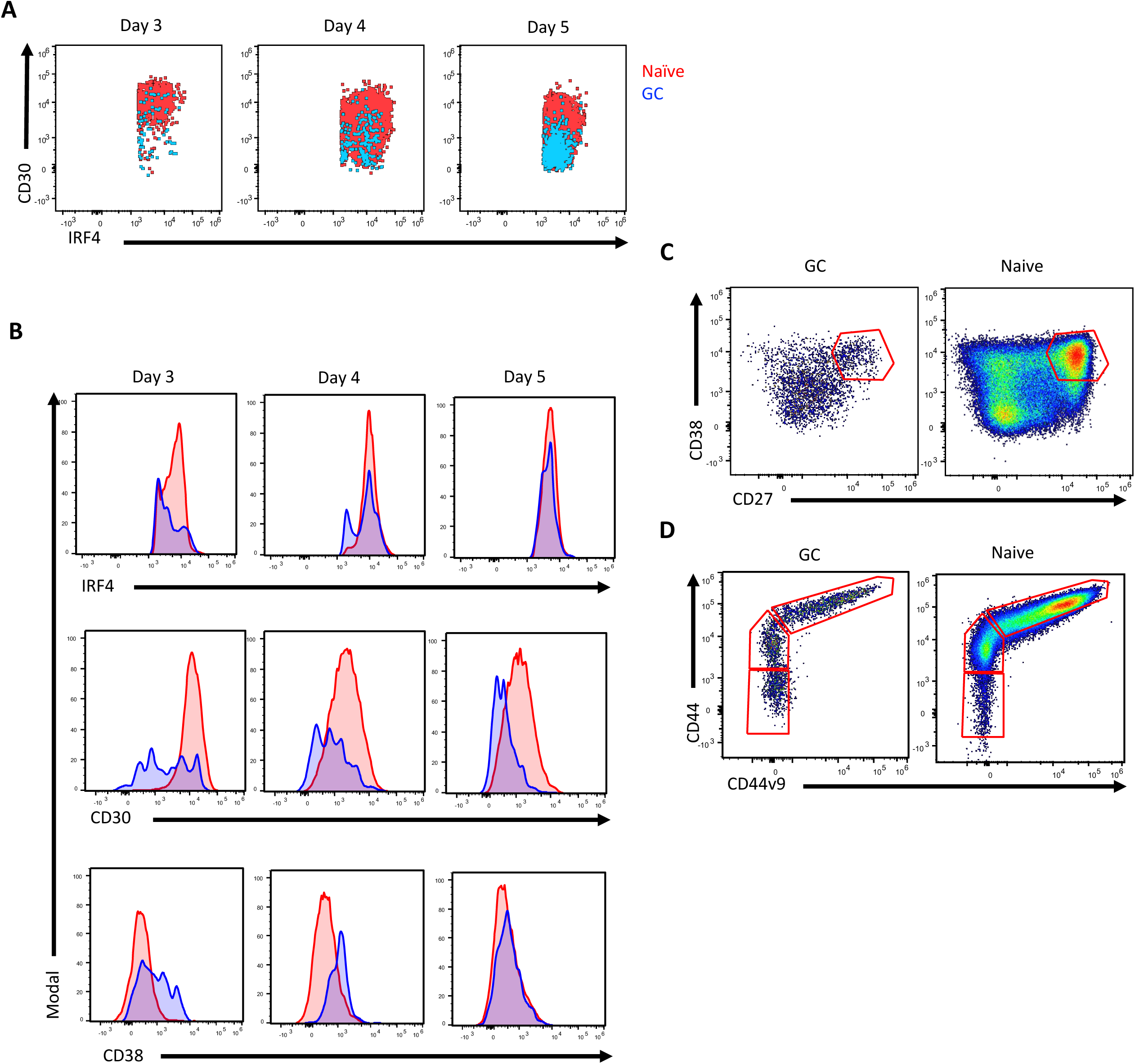
Primary germinal center B cells fail to form the CD30- and IRF4-based populations observed with naïve B cell. **(A)** Daily time course of CD30 and IRF4 expression for *in vitro* differentiating sorted naïve B cells or germinal center B cells showing only those with IRF4 expression above the level of unresponsive cells (best visualized in Fig 4B day 3). **(B)** Overlaid histograms for the highest IRF4 expressing cells looking at IRF4, CD30, and CD38 expression. **(C)** Representative CD27 and CD38 profiles at the end of differentiation starting from germinal center or naïve B cells. **(D)** Representative CD44 and CD44v9 profiles at the end of differentiation starting from germinal center or naïve B cells.

**Sup. Fig. 5:**
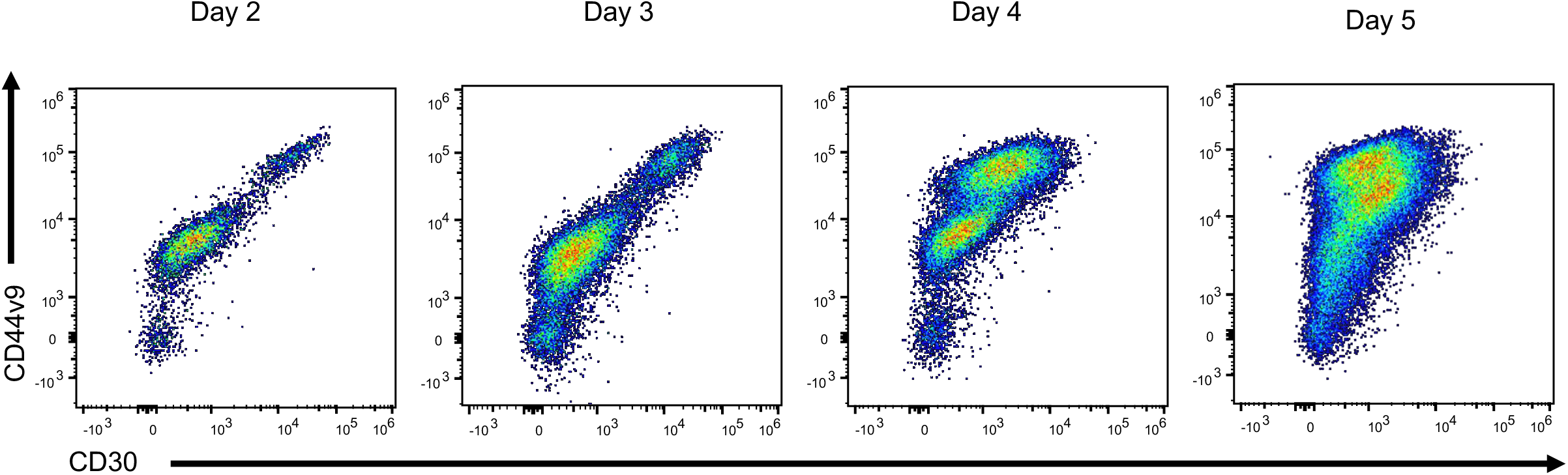
Validation of linked CD30 and CD44v9 expression. Daily time course of CD30 and CD44v9 expression of *in vitro* differentiating naïve B cells.

**Sup. Fig. 6:**
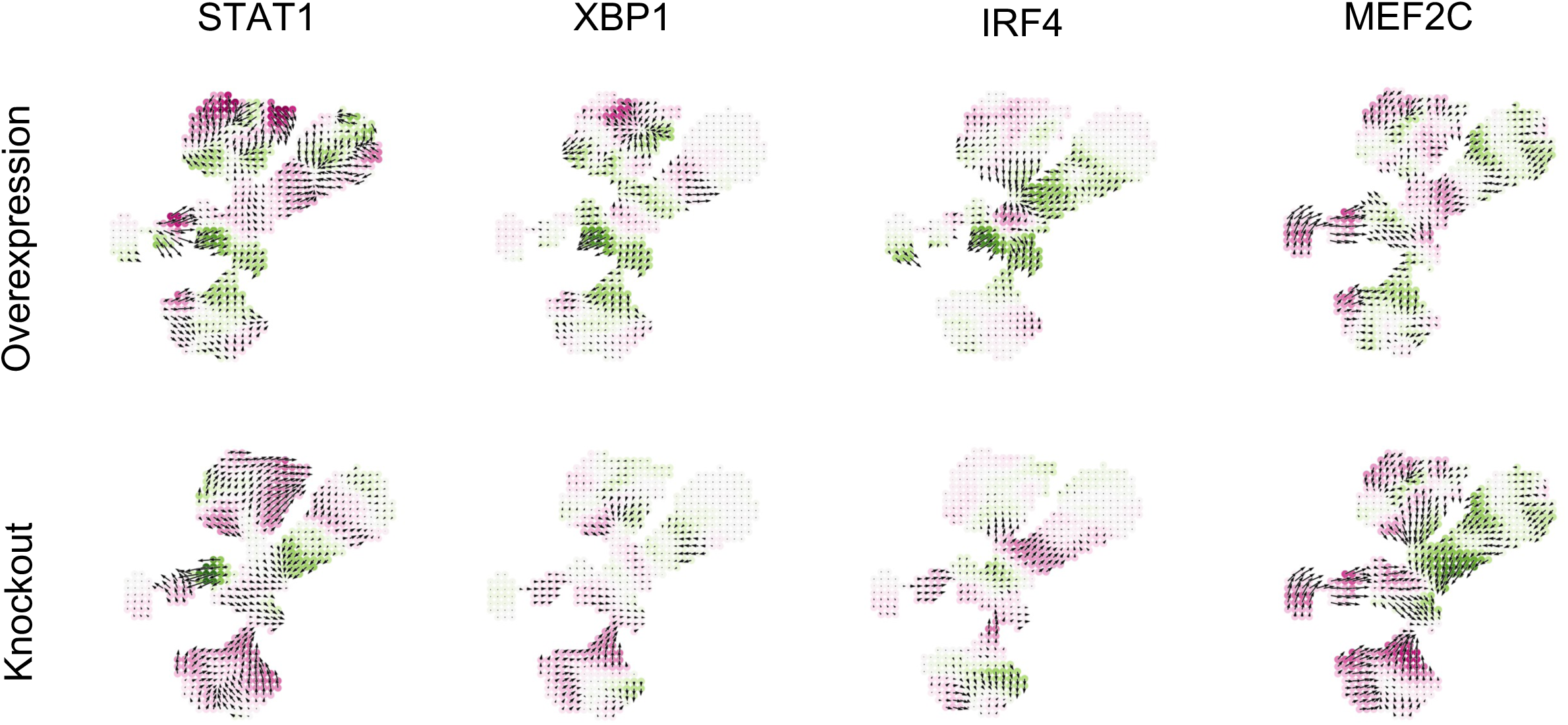
Simulated overexpression or knockout of transcription factors identified by CellOracle analysis. Overexpression was assessed by a simulated 10 to 20% increase in transcription factor activity and complete knockout for STAT1, XBP1 (as a surrogate for rosiglitazone activity), IRF4, and MEF2C. Green coloration indicates changes predicted to progress towards the plasma cell identity while purple coloration indicates predicted changes away from the plasma cell identity.

**Sup. Fig. 7:**
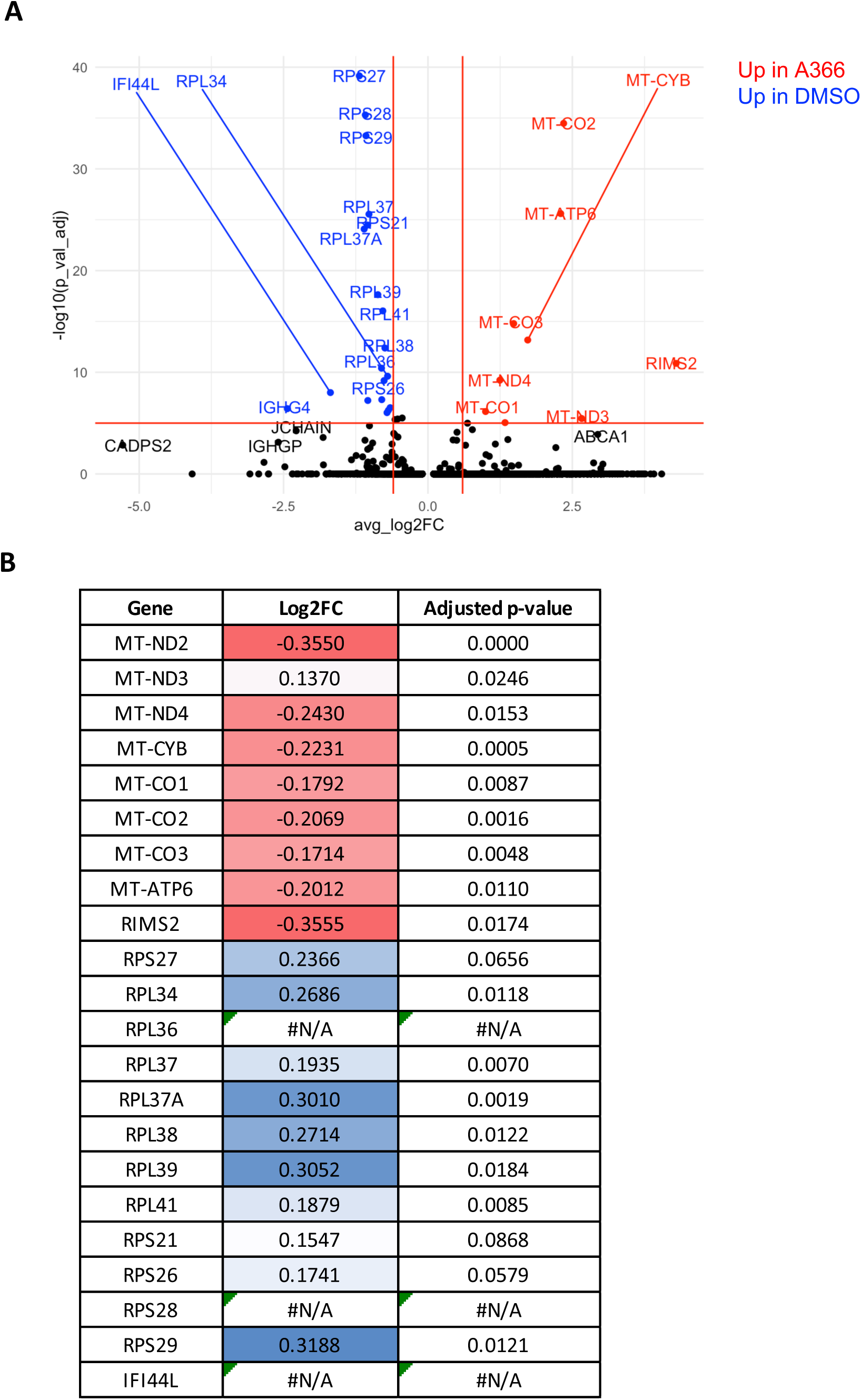
Differentially expressed genes in A366 treated cells match with MEF2C direct targets. **(A)** Differentially expressed genes between A366 and DMSO treated cells. **(B)** Log-2-fold change and adjusted p-values for the genes identified in **(A)** comparing MEF2C knockout cells to wildtype cells reported in Ref. (Ow et al., 2016).

**Sup. Fig. 8:**
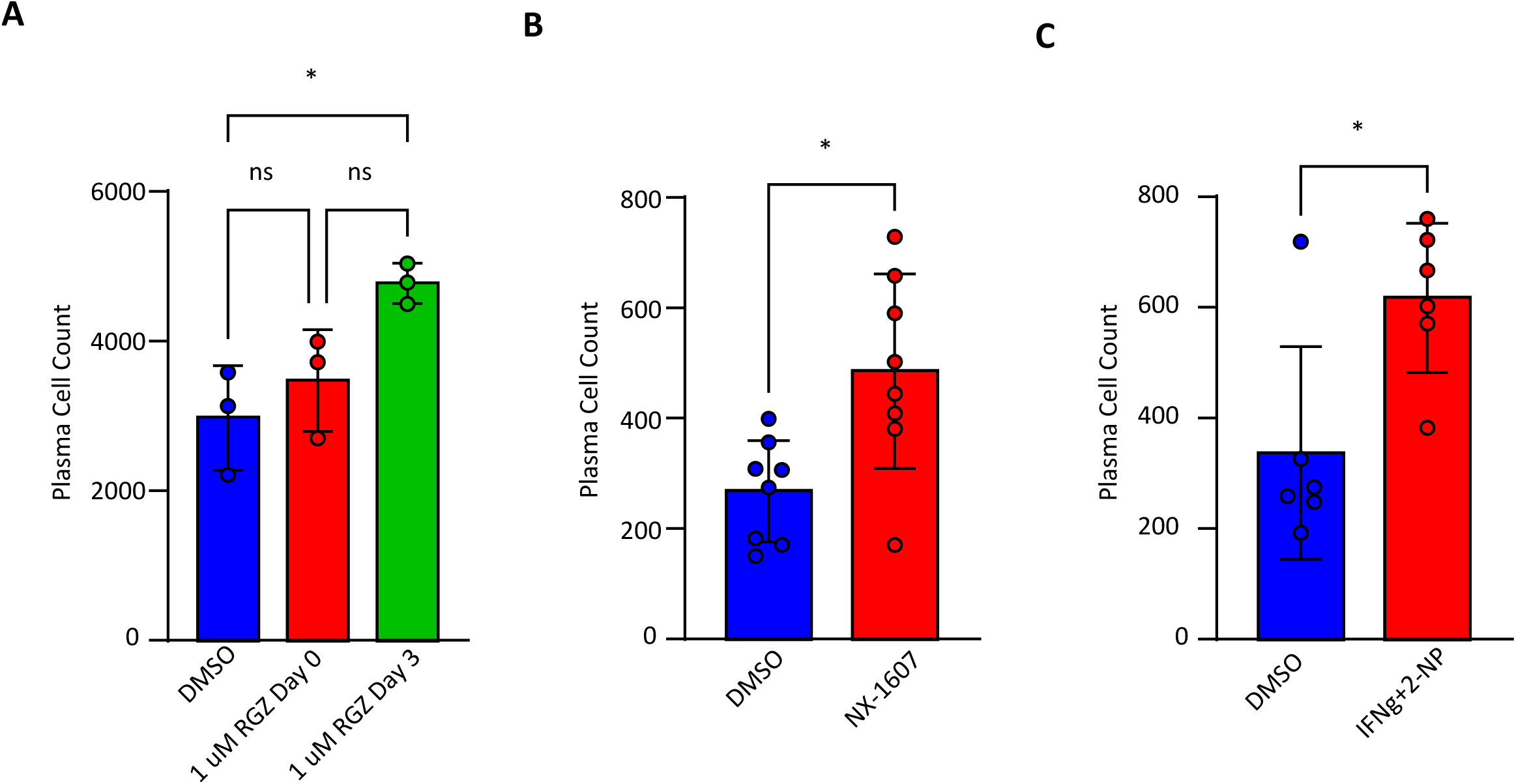
Increased Xbp1, IRF4, and STAT signaling improve plasma cell yields. Final plasma cell yields for **(A)** Rosiglitazone added on Day 0 or 3, **(B)** NX-1607 added on Day 5, or **(C)** Interferon-γ and 2-NP added on Day 5. Statistical significance was evaluated using a t-test in figures where only two groups are being compared. In all other instances, one-way analysis of variance (ANOVA) was performed to test for statistical significance. Data are presented as mean ± standard error of the mean (SEM). Statistical significance was defined as *p < 0.05.

**Sup. Fig. 9:**
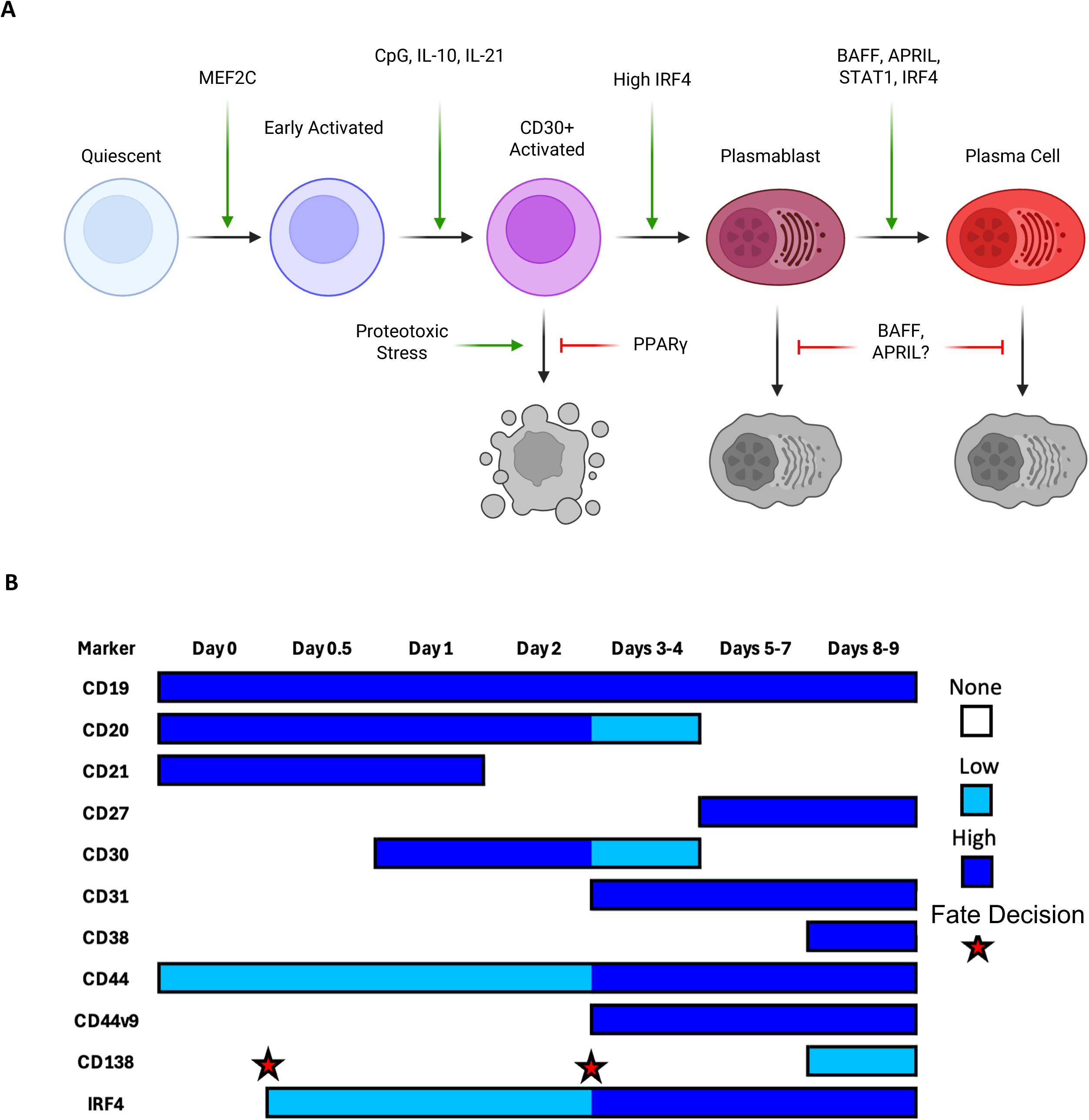
Regulation of plasma cell differentiation and characteristic changes observed during differentiation. **(A)** Identified and functionally validated regulators of different stages of plasma cell differentiation. **(B)** Changes in protein expression observed during *in vitro* plasma cell differentiation. Potential fate decisions are indicated with a red star.

## References

Alaterre, E., Ovejero, S., Bret, C., Dutrieux, L., Sika, D., Fernandez Perez, R., Espéli, M., Fest, T., Cogné, M., Martin-Subero, J.I., Milpied, P., Cavalli, G., Moreaux, J., 2024. Integrative single-cell chromatin and transcriptome analysis of human plasma cell differentiation. Blood 144, 496–509. 10.1182/blood.2023023237

Alquicira-Hernandez, J., Powell, J.E., Phan, T.G., 2021. No evidence that plasmablasts transdifferentiate into developing neutrophils in severe COVID-19 disease. Clinical & Translational Immunology 10, e1308. 10.1002/cti2.1308

Amanna, I.J., Carlson, N.E., Slifka, M.K., 2007. Duration of Humoral Immunity to Common Viral and Vaccine Antigens. New England Journal of Medicine 357, 1903–1915. 10.1056/NEJMoa066092

Bekeredjian-Ding, I., Jego, G., 2009. Toll-like receptors – sentries in the B-cell response. Immunology 128, 311–323. 10.1111/j.1365-2567.2009.03173.x

Bortnick, A., Allman, D., 2013. What Is and What Should Always Have Been: Long-Lived Plasma Cells Induced by T Cell–Independent Antigens. J Immunol 190, 5913–5918. 10.4049/jimmunol.1300161

Brynjolfsson, S.F., Mohaddes, M., Kärrholm, J., Wick, M.-J., 2017. Long-lived plasma cells in human bone marrow can be either CD19+ or CD19–. Blood Advances 1, 835–838. 10.1182/bloodadvances.2017004481

Calderón, L., Schäfer, M., Rončević, M., Rauschmeier, R., Jaritz, M., Schwickert, T.A., Sun, Q., Pauli, A., Zuber, J., Busslinger, M., 2026. In vivo CRISPR/Cas9 screens identify new regulators of B cell activation and plasma cell differentiation. J Exp Med 223, e20250594. 10.1084/jem.20250594

Cao, J., Spielmann, M., Qiu, X., Huang, X., Ibrahim, D.M., Hill, A.J., Zhang, F., Mundlos, S., Christiansen, L., Steemers, F.J., Trapnell, C., Shendure, J., 2019. The single-cell transcriptional landscape of mammalian organogenesis. Nature 566, 496–502. 10.1038/s41586-019-0969-x

Chari, T., Pachter, L., 2023. The specious art of single-cell genomics. PLOS Computational Biology 19, e1011288. 10.1371/journal.pcbi.1011288

Cheekati, M., Murakhovskaya, I., 2024. Anti-B-Cell-Activating Factor (BAFF) Therapy: A Novel Addition to Autoimmune Disease Management and Potential for Immunomodulatory Therapy in Warm Autoimmune Hemolytic Anemia. Biomedicines 12, 1597. 10.3390/biomedicines12071597

Debnath, I., Roundy, K.M., Pioli, P.D., Weis, J.J., Weis, J.H., 2013. Bone marrow-induced Mef2c deficiency delays B-cell development and alters the expression of key B-cell regulatory proteins. Int Immunol 25, 99–115. 10.1093/intimm/dxs088

DiLillo, D.J., Hamaguchi, Y., Ueda, Y., Yang, K., Uchida, J., Haas, K.M., Kelsoe, G., Tedder, T.F., 2008. Maintenance of Long-Lived Plasma Cells and Serological Memory Despite Mature and Memory B Cell Depletion during CD20 Immunotherapy in Mice1. The Journal of Immunology 180, 361–371. 10.4049/jimmunol.180.1.361

Eisenbarth, S.C., Batista, F., Cyster, J., Elsner, R., Kelsoe, G., Lund, F.E., Pillai, S., Sanz, I., Shlomchik, M., Toellner, K.-M., Vinuesa, C., Baumgarth, N., 2025. A roadmap for defining “extrafollicular” B cell responses. Immunity 58, 2627–2645. 10.1016/j.immuni.2025.08.007

Ferreira-Gomes, M., Chen, Y., Durek, P., Rincon-Arevalo, H., Heinrich, F., Bauer, L., Szelinski, F., Guerra, G.M., Stefanski, A.-L., Niedobitek, A., Wiedemann, A., Bondareva, M., Ritter, J., Lehmann, K., Hardt, S., Hipfl, C., Hein, S., Hildt, E., Matz, M., Mei, H.E., Cheng, Q., Dang, V.D., Witkowski, M., Lino, A.C., Kruglov, A., Melchers, F., Perka, C., Schrezenmeier, E.V., Hutloff, A., Radbruch, A., Dörner, T., Mashreghi, M.-F., 2024. Recruitment of plasma cells from IL-21-dependent and IL-21-independent immune reactions to the bone marrow. Nat Commun 15, 4182. 10.1038/s41467-024-48570-0

Fritz, J.H., Rojas, O.L., Simard, N., McCarthy, D.D., Hapfelmeier, S., Rubino, S., Robertson, S.J., Larijani, M., Gosselin, J., Ivanov, I.I., Martin, A., Casellas, R., Philpott, D.J., Girardin, S.E., McCoy, K.D., Macpherson, A.J., Paige, C.J., Gommerman, J.L., 2012. Acquisition of a multifunctional IgA+ plasma cell phenotype in the gut. Nature 481, 199–203. 10.1038/nature10698

Gallotta, M., Gosling, J., Tenn-McClellan, A., Ranucci, S., Romo, J.G., Cohen, F., Hansen, G., Sands, A., Guiducci, C., Rountree, R., 2022. 824 NX-1607, a small molecule inhibitor of the CBL-B E3 ubiquitin ligase, promotes T and NK cell activation and enhances NK-mediated ADCC in a mouse lymphoma tumor model. J Immunother Cancer 10. 10.1136/jitc-2022-SITC2022.0824

Garcia-Bates, T.M., Baglole, C.J., Bernard, M.P., Murant, T.I., Simpson-Haidaris, P.J., Phipps, R.P., 2009. Peroxisome Proliferator-Activated Receptor γ Ligands Enhance Human B Cell Antibody Production and Differentiation1. J Immunol 183, 6903–6912. 10.4049/jimmunol.0900324

Gass, J.N., Gifford, N.M., Brewer, J.W., 2002. Activation of an Unfolded Protein Response during Differentiation of Antibody-secreting B Cells*. Journal of Biological Chemistry 277, 49047–49054. 10.1074/jbc.M205011200

Gaudette, B.T., Jones, D.D., Bortnick, A., Argon, Y., Allman, D., 2020. mTORC1 coordinates an immediate unfolded protein response-related transcriptome in activated B cells preceding antibody secretion. Nat Commun 11, 723. 10.1038/s41467-019-14032-1

Gómez-Escolar, C., Serrano-Navarro, A., Benguria, A., Dopazo, A., Sánchez-Cabo, F., Ramiro, A.R., 2022. Single cell clonal analysis identifies an AID-dependent pathway of plasma cell differentiation. EMBO Rep 23, EMBR202255000. 10.15252/embr.202255000

Halliley, J.L., Tipton, C.M., Liesveld, J., Rosenberg, A.F., Darce, J., Gregoretti, I.V., Popova, L., Kaminiski, D., Fucile, C.F., Albizua, I., Kyu, S., Chiang, K.-Y., Bradley, K.T., Burack, R., Slifka, M., Hammarlund, E., Wu, H., Zhao, L., Walsh, E.E., Falsey, A.R., Randall, T.D., Cheung, W.C., Sanz, I., Lee, F.E.-H., 2015. Long-Lived Plasma Cells Are Contained within the CD19−CD38hiCD138+ Subset in Human Bone Marrow. Immunity 43, 132–145. 10.1016/j.immuni.2015.06.016

Higashioka, K., Kikushige, Y., Ayano, M., Kimoto, Y., Mitoma, H., Kikukawa, M., Akahoshi, M., Arinobu, Y., Horiuchi, T., Akashi, K., Niiro, H., 2020. Generation of a novel CD30+ B cell subset producing GM-CSF and its possible link to the pathogenesis of systemic sclerosis. Clin Exp Immunol 201, 233–243. 10.1111/cei.13477

Higgins, B.W., Shuparski, A.G., Miller, K.B., Robinson, A.M., McHeyzer-Williams, L.J., McHeyzer-Williams, M.G., 2022. Isotype-specific plasma cells express divergent transcriptional programs. Proceedings of the National Academy of Sciences 119, e2121260119. 10.1073/pnas.2121260119

Hipp, N., Symington, H., Pastoret, C., Caron, G., Monvoisin, C., Tarte, K., Fest, T., Delaloy, C., 2017. IL-2 imprints human naive B cell fate towards plasma cell through ERK/ELK1-mediated BACH2 repression. Nat Commun 8, 1443. 10.1038/s41467-017-01475-7

Hwang, I.-Y., Park, C., Harrison, K., Kehrl, J.H., 2009. TLR4 signaling augments B lymphocyte migration and overcomes the restriction that limits access to germinal center dark zones. J Exp Med 206, 2641–2657. 10.1084/jem.20091982

Ise, W., Fujii, K., Shiroguchi, K., Ito, A., Kometani, K., Takeda, K., Kawakami, E., Yamashita, K., Suzuki, K., Okada, T., Kurosaki, T., 2018. T Follicular Helper Cell-Germinal Center B Cell Interaction Strength Regulates Entry into Plasma Cell or Recycling Germinal Center Cell Fate. Immunity 48, 702–715.e4. 10.1016/j.immuni.2018.03.027

Ise, W., Koike, T., Shimada, N., Yamamoto, H., Tai, Y., Shirai, T., Kawakami, R., Kuwabara, M., Kawai, C., Shida, K., Inoue, T., Hojo, N., Ichiyama, K., Sakaguchi, S., Shiroguchi, K., Suzuki, K., Kurosaki, T., 2025. KLF2 expression in IgG plasma cells at their induction site regulates the migration program. J Exp Med 222, e20241019. 10.1084/jem.20241019

Jego, G., Palucka, A.K., Blanck, J.-P., Chalouni, C., Pascual, V., Banchereau, J., 2003. Plasmacytoid Dendritic Cells Induce Plasma Cell Differentiation through Type I Interferon and Interleukin 6. Immunity 19, 225–234. 10.1016/S1074-7613(03)00208-5

Jourdan, M., Caraux, A., De Vos, J., Fiol, G., Larroque, M., Cognot, C., Bret, C., Duperray, C., Hose, D., Klein, B., 2009. An in vitro model of differentiation of memory B cells into plasmablasts and plasma cells including detailed phenotypic and molecular characterization. Blood 114, 5173–5181. 10.1182/blood-2009-07-235960

Jourdan, M., Cren, M., Robert, N., Bolloré, K., Fest, T., Duperray, C., Guilloton, F., Hose, D., Tarte, K., Klein, B., 2014. IL-6 supports the generation of human long-lived plasma cells in combination with either APRIL or stromal cell-soluble factors. Leukemia 28, 1647–1656. 10.1038/leu.2014.61

Kamimoto, K., Stringa, B., Hoffmann, C.M., Jindal, K., Solnica-Krezel, L., Morris, S.A., 2023. Dissecting cell identity via network inference and in silico gene perturbation. Nature 614, 742–751. 10.1038/s41586-022-05688-9

Kim, S.V., Xiang, W.V., Kwak, C., Yang, Y., Lin, X.W., Ota, M., Sarpel, U., Rifkin, D.B., Xu, R., Littman, D.R., 2013. GPR15-Mediated Homing Controls Immune Homeostasis in the Large Intestine Mucosa. Science 340, 1456–1459. 10.1126/science.1237013

Kräutler, N.J., Suan, D., Butt, D., Bourne, K., Hermes, J.R., Chan, T.D., Sundling, C., Kaplan, W., Schofield, P., Jackson, J., Basten, A., Christ, D., Brink, R., 2017a. Differentiation of germinal center B cells into plasma cells is initiated by high-affinity antigen and completed by Tfh cells. J Exp Med 214, 1259–1267. 10.1084/jem.20161533

Kräutler, N.J., Suan, D., Butt, D., Bourne, K., Hermes, J.R., Chan, T.D., Sundling, C., Kaplan, W., Schofield, P., Jackson, J., Basten, A., Christ, D., Brink, R., 2017b. Differentiation of germinal center B cells into plasma cells is initiated by high-affinity antigen and completed by Tfh cells. J Exp Med 214, 1259–1267. 10.1084/jem.20161533

Lam, W.Y., Jash, A., Yao, C.-H., D’Souza, L., Wong, R., Nunley, R.M., Meares, G.P., Patti, G.J., Bhattacharya, D., 2018. Metabolic and Transcriptional Modules Independently Diversify Plasma Cell Lifespan and Function. Cell Reports 24, 2479–2492.e6. 10.1016/j.celrep.2018.07.084

Landsverk, O.J.B., Snir, O., Casado, R.B., Richter, L., Mold, J.E., Réu, P., Horneland, R., Paulsen, V., Yaqub, S., Aandahl, E.M., Øyen, O.M., Thorarensen, H.S., Salehpour, M., Possnert, G., Frisén, J., Sollid, L.M., Baekkevold, E.S., Jahnsen, F.L., 2017. Antibody-secreting plasma cells persist for decades in human intestine. J Exp Med 214, 309–317. 10.1084/jem.20161590

Lemke, A., Kraft, M., Roth, K., Riedel, R., Lammerding, D., Hauser, A.E., 2016. Long-lived plasma cells are generated in mucosal immune responses and contribute to the bone marrow plasma cell pool in mice. Mucosal Immunol 9, 83–97. 10.1038/mi.2015.38

Li, X., Gadzinsky, A., Gong, L., Tong, H., Calderon, V., Li, Y., Kitamura, D., Klein, U., Langdon, W.Y., Hou, F., Zou, Y.-R., Gu, H., 2018. Cbl Ubiquitin Ligases Control B Cell Exit from the Germinal-Center Reaction. Immunity 48, 530–541.e6. 10.1016/j.immuni.2018.03.006

Lin, K.-I., Angelin-Duclos, C., Kuo, T.C., Calame, K., 2002. Blimp-1-Dependent Repression of Pax-5 Is Required for Differentiation of B Cells to Immunoglobulin M-Secreting Plasma Cells. Molecular and Cellular Biology 22, 4771–4780. 10.1128/MCB.22.13.4771-4780.2002

Lynch, R.A., Etchin, J., Battle, T.E., Frank, D.A., 2007. A Small-Molecule Enhancer of Signal Transducer and Activator of Transcription 1 Transcriptional Activity Accentuates the Antiproliferative Effects of IFN-γ in Human Cancer Cells. Cancer Res 67, 1254–1261. 10.1158/0008-5472.CAN-06-2439

Matsuda, Y., Haneda, M., Kadomatsu, K., Kobayashi, T., 2015. A proliferation-inducing ligand sustains the proliferation of human naïve (CD27−) B cells and mediates their differentiation into long-lived plasma cells *in vitro* via transmembrane activator and calcium modulator and cyclophilin ligand interactor and B-cell mature antigen. Cellular Immunology 295, 127–136. 10.1016/j.cellimm.2015.02.011

Mei, H.E., Wirries, I., Frölich, D., Brisslert, M., Giesecke, C., Grün, J.R., Alexander, T., Schmidt, S., Luda, K., Kühl, A.A., Engelmann, R., Dürr, M., Scheel, T., Bokarewa, M., Perka, C., Radbruch, A., Dörner, T., 2015. A unique population of IgG-expressing plasma cells lacking CD19 is enriched in human bone marrow. Blood 125, 1739–1748. 10.1182/blood-2014-02-555169

Meng, Y., Ying, M., Hongwen, Z., Li, H., Xiaopeng, T., 2025. Telitacicept for lupus nephritis with BAFF and APRIL double positivity in children: a case report. BMC Pediatr 25, 422. 10.1186/s12887-025-05778-3

Mesin, L., Schiepers, A., Ersching, J., Barbulescu, A., Cavazzoni, C.B., Angelini, A., Okada, T., Kurosaki, T., Victora, G.D., 2020. Restricted Clonality and Limited Germinal Center Reentry Characterize Memory B Cell Reactivation by Boosting. Cell 180, 92–106.e11. 10.1016/j.cell.2019.11.032

Mosser, D.M., Edwards, J.P., 2008. Exploring the full spectrum of macrophage activation. Nat Rev Immunol 8, 958–969. 10.1038/nri2448

Murray, P.J., Allen, J.E., Biswas, S.K., Fisher, E.A., Gilroy, D.W., Goerdt, S., Gordon, S., Hamilton, J.A., Ivashkiv, L.B., Lawrence, T., Locati, M., Mantovani, A., Martinez, F.O., Mege, J.-L., Mosser, D.M., Natoli, G., Saeij, J.P., Schultze, J.L., Shirey, K.A., Sica, A., Suttles, J., Udalova, I., van Ginderachter, J.A., Vogel, S.N., Wynn, T.A., 2014. Macrophage Activation and Polarization: Nomenclature and Experimental Guidelines. Immunity 41, 14–20. 10.1016/j.immuni.2014.06.008

Orecchioni, M., Ghosheh, Y., Pramod, A.B., Ley, K., 2019. Macrophage Polarization: Different Gene Signatures in M1(LPS+) vs. Classically and M2(LPS–) vs. Alternatively Activated Macrophages. Front. Immunol. 10. 10.3389/fimmu.2019.01084

Ow, J.R., Palanichamy Kala, M., Rao, V.K., Choi, M.H., Bharathy, N., Taneja, R., 2016. G9a inhibits MEF2C activity to control sarcomere assembly. Sci Rep 6, 34163. 10.1038/srep34163

Poeck, H., Wagner, M., Battiany, J., Rothenfusser, S., Wellisch, D., Hornung, V., Jahrsdorfer, B., Giese, T., Endres, S., Hartmann, G., 2004. Plasmacytoid dendritic cells, antigen, and CpG-C license human B cells for plasma cell differentiation and immunoglobulin production in the absence of T-cell help. Blood 103, 3058–3064. 10.1182/blood-2003-08-2972

Salmi, M., Jalkanen, S., 2005. Cell-surface enzymes in control of leukocyte trafficking. Nat Rev Immunol 5, 760–771. 10.1038/nri1705

Schiepers, A., van ’t Wout, M.F.L., Greaney, A.J., Zang, T., Muramatsu, H., Lin, P.J.C., Tam, Y.K., Mesin, L., Starr, T.N., Bieniasz, P.D., Pardi, N., Bloom, J.D., Victora, G.D., 2023. Molecular fate-mapping of serum antibody responses to repeat immunization. Nature 615, 482–489. 10.1038/s41586-023-05715-3

Schulz, S.R., Menzel, S.R., Wittner, J., Ulbricht, C., Grofe, A.T., Roth, E., Mann-Nüttel, R., Scheu, S., Kueh, A.J., Jäck, A., Herold, M.J., Hauser, A.E., Pracht, K., Schuh, W., Jäck, H.-M., 2025. Decoding plasma cell maturation dynamics with BCMA. Front. Immunol. 16. 10.3389/fimmu.2025.1539773

Sciammas, R., Shaffer, A.L., Schatz, J.H., Zhao, H., Staudt, L.M., Singh, H., 2006. Graded Expression of Interferon Regulatory Factor-4 Coordinates Isotype Switching with Plasma Cell Differentiation. Immunity 25, 225–236. 10.1016/j.immuni.2006.07.009

Tellier, J., Tarasova, I., Nie, J., Smillie, C.S., Fedele, P.L., Cao, W.H.J., Groom, J.R., Belz, G.T., Bhattacharya, D., Smyth, G.K., Nutt, S.L., 2024a. Unraveling the diversity and functions of tissue-resident plasma cells. Nat Immunol 25, 330–342. 10.1038/s41590-023-01712-w

Tellier, J., Tarasova, I., Nie, J., Smillie, C.S., Fedele, P.L., Cao, W.H.J., Groom, J.R., Belz, G.T., Bhattacharya, D., Smyth, G.K., Nutt, S.L., 2024b. Unraveling the diversity and functions of tissue-resident plasma cells. Nat Immunol 25, 330–342. 10.1038/s41590-023-01712-w

Tian, Z., Yang, Y., Huang, M., Fang, Z., Mei, J., Li, Yunyi, Tang, L., Li, Yanyan, Li, Yuxia, 2025. Efficacy and safety of BAFF/APRIL dual antagonists in IgA nephropathy: a systematic review and meta-analysis of randomized controlled trials. BMC Nephrol 27, 55. 10.1186/s12882-025-04689-w

Tsui, C., Martinez-Martin, N., Gaya, M., Maldonado, P., Llorian, M., Legrave, N.M., Rossi, M., MacRae, J.I., Cameron, A.J., Parker, P.J., Leitges, M., Bruckbauer, A., Batista, F.D., 2018. Protein Kinase C-β Dictates B Cell Fate by Regulating Mitochondrial Remodeling, Metabolic Reprogramming, and Heme Biosynthesis. Immunity 48, 1144–1159.e5. 10.1016/j.immuni.2018.04.031

Turner, J.S., Zhou, J.Q., Han, J., Schmitz, A.J., Rizk, A.A., Alsoussi, W.B., Lei, T., Amor, M., McIntire, K.M., Meade, P., Strohmeier, S., Brent, R.I., Richey, S.T., Haile, A., Yang, Y.R., Klebert, M.K., Suessen, T., Teefey, S., Presti, R.M., Krammer, F., Kleinstein, S.H., Ward, A.B., Ellebedy, A.H., 2020. Human germinal centres engage memory and naive B cells after influenza vaccination. Nature 586, 127–132. 10.1038/s41586-020-2711-0

Van Driel, M., Günthert, U., van Kessel, A.C., Joling, P., Stauder, R., Lokhorst, H.M., Bloem, A.C., 2002. CD44 variant isoforms are involved in plasma cell adhesion to bone marrow stromal cells. Leukemia 16, 135–143. 10.1038/sj.leu.2402336

van Spriel, A.B., de Keijzer, S., van der Schaaf, A., Gartlan, K.H., Sofi, M., Light, A., Linssen, P.C., Boezeman, J.B., Zuidscherwoude, M., Reinieren-Beeren, I., Cambi, A., Mackay, F., Tarlinton, D.M., Figdor, C.G., Wright, M.D., 2012. The Tetraspanin CD37 Orchestrates the α4β1 Integrin–Akt Signaling Axis and Supports Long-Lived Plasma Cell Survival. Science Signaling 5, ra82–ra82. 10.1126/scisignal.2003113

Vecchione, A., Devlin, J.C., Tasker, C., Ramnarayan, V.R., Haase, P., Conde, E., Srivastava, D., Atwal, G.S., Bruhns, P., Murphy, A.J., Sleeman, M.A., Limnander, A., Lim, W.K., Asrat, S., Orengo, J.M., 2024. IgE plasma cells are transcriptionally and functionally distinct from other isotypes. Science Immunology 9, eadm8964. 10.1126/sciimmunol.adm8964

Verstegen, N.J., Pollastro, S., Unger, P.-P.A., Marsman, C., Elias, G., Jorritsma, T., Streutker, M., Bassler, K., Haendler, K., Rispens, T., Schultze, J.L., ten Brinke, A., Beyer, M., van Ham, S.M., 2023. Single-cell analysis reveals dynamics of human B cell differentiation and identifies novel B and antibody-secreting cell intermediates. eLife 12, e83578. 10.7554/eLife.83578

Wang, S., Wang, J., Kumar, V., Karnell, J.L., Naiman, B., Gross, P.S., Rahman, S., Zerrouki, K., Hanna, R., Morehouse, C., Holoweckyj, N., Liu, H., Manna, Z., Goldbach-Mansky, R., Hasni, S., Siegel, R., Sanjuan, M., Streicher, K., Cancro, M.P., Kolbeck, R., Ettinger, R., 2018. IL-21 drives expansion and plasma cell differentiation of autoreactive CD11chiT-bet+ B cells in SLE. Nat Commun 9, 1758. 10.1038/s41467-018-03750-7

Weniger, M.A., Tiacci, E., Schneider, S., Arnolds, J., Rüschenbaum, S., Duppach, J., Seifert, M., Döring, C., Hansmann, M.-L., Küppers, R., 2018. Human CD30^+^ B cells represent a unique subset related to Hodgkin lymphoma cells. J Clin Invest 128, 2996–3007. 10.1172/JCI95993

White, M.T., Verity, R., Griffin, J.T., Asante, K.P., Owusu-Agyei, S., Greenwood, B., Drakeley, C., Gesase, S., Lusingu, J., Ansong, D., Adjei, S., Agbenyega, T., Ogutu, B., Otieno, L., Otieno, W., Agnandji, S.T., Lell, B., Kremsner, P., Hoffman, I., Martinson, F., Kamthunzu, P., Tinto, H., Valea, I., Sorgho, H., Oneko, M., Otieno, K., Hamel, M.J., Salim, N., Mtoro, A., Abdulla, S., Aide, P., Sacarlal, J., Aponte, J.J., Njuguna, P., Marsh, K., Bejon, P., Riley, E.M., Ghani, A.C., 2015. Immunogenicity of the RTS,S/AS01 malaria vaccine and implications for duration of vaccine efficacy: secondary analysis of data from a phase 3 randomised controlled trial. The Lancet Infectious Diseases 15, 1450–1458. 10.1016/S1473-3099(15)00239-X

Wilk, A.J., Rustagi, A., Zhao, N.Q., Roque, J., Martínez-Colón, G.J., McKechnie, J.L., Ivison, G.T., Ranganath, T., Vergara, R., Hollis, T., Simpson, L.J., Grant, P., Subramanian, A., Rogers, A.J., Blish, C.A., 2020. A single-cell atlas of the peripheral immune response in patients with severe COVID-19. Nat Med 26, 1070–1076. 10.1038/s41591-020-0944-y

Wilker, P.R., Kohyama, M., Sandau, M.M., Albring, J.C., Nakagawa, O., Schwarz, J.J., Murphy, K.M., 2008. Transcription factor Mef2c is required for B cell proliferation and survival after antigen receptor stimulation. Nat Immunol 9, 603–612. 10.1038/ni.1609

Wilmore, J.R., Gaudette, B.T., Gomez Atria, D., Hashemi, T., Jones, D.D., Gardner, C.A., Cole, S.D., Misic, A.M., Beiting, D.P., Allman, D., 2018. Commensal Microbes Induce Serum IgA Responses that Protect against Polymicrobial Sepsis. Cell Host & Microbe 23, 302–311.e3. 10.1016/j.chom.2018.01.005

Wilmore, J.R., Gaudette, B.T., Gómez Atria, D., Rosenthal, R.L., Reiser, S.K., Meng, W., Rosenfeld, A.M., Luning Prak, E.T., Allman, D., 2021. IgA Plasma Cells Are Long-Lived Residents of Gut and Bone Marrow That Express Isotype- and Tissue-Specific Gene Expression Patterns. Front. Immunol. 12. 10.3389/fimmu.2021.791095

Wols, H.A.M., Underhill, G.H., Kansas, G.S., Witte, P.L., 2002. The Role of Bone Marrow-Derived Stromal Cells in the Maintenance of Plasma Cell Longevity1. J Immunol 169, 4213–4221. 10.4049/jimmunol.169.8.4213

Wong, R., Belk, J.A., Govero, J., Uhrlaub, J.L., Reinartz, D., Zhao, H., Errico, J.M., D’Souza, L., Ripperger, T.J., Nikolich-Zugich, J., Shlomchik, M.J., Satpathy, A.T., Fremont, D.H., Diamond, M.S., Bhattacharya, D., 2020. Affinity-Restricted Memory B Cells Dominate Recall Responses to Heterologous Flaviviruses. Immunity 53, 1078–1094.e7. 10.1016/j.immuni.2020.09.001

Xie, Z., Geiger, T.R., Johnson, E.N., Nyborg, J.K., Druey, K.M., 2008. RGS13 Acts as a Nuclear Repressor of CREB. Molecular Cell 31, 660–670. 10.1016/j.molcel.2008.06.024

Young, C., Brink, R., 2021. The unique biology of germinal center B cells. Immunity 54, 1652–1664. 10.1016/j.immuni.2021.07.015

Zumaquero, E., Stone, S.L., Scharer, C.D., Jenks, S.A., Nellore, A., Mousseau, B., Rosal-Vela, A., Botta, D., Bradley, J.E., Wojciechowski, W., Ptacek, T., Danila, M.I., Edberg, J.C., Bridges, S.L., Jr, Kimberly, R.P., Chatham, W.W., Schoeb, T.R., Rosenberg, A.F., Boss, J.M., Sanz, I., Lund, F.E., 2019. IFNγ induces epigenetic programming of human T-bethi B cells and promotes TLR7/8 and IL-21 induced differentiation. eLife 8, e41641. 10.7554/eLife.41641

